# Comprehensive characterization of the integrin family across 32 cancer types

**DOI:** 10.1101/2023.12.24.573245

**Authors:** Cheng Zou, Jinwei Zhu, Jiangling Xiong, Yu Tian, Yousong Peng, Edwin Cheung, Dingxiao Zhang

## Abstract

Integrin genes widely involve in tumorigenesis. Yet, a comprehensive characterization of integrin family and their interactome on the pan-cancer level is lacking. Here, we systematically dissect integrin family in nearly 10000 tumors across 32 cancer types. Globally, integrins represent a frequently altered and misexpressed pathway, with alteration and dysregulation overall being protumorigenic. Expression dysregulation, better than mutational landscape, of integrin family successfully identifies a subgroup of aggressive tumors demonstrating a high level of proliferation and stemness. We identify that several molecular mechanisms jointly regulate integrin expression in a context-dependent manner. For potential clinical usage, we construct a weighted score, integrinScore, to measure integrin signaling patterns in individual tumors. Remarkably, integrinScore consistently correlates with predefined molecular subtypes in multiple cancers, with integrinScore high tumors being more aggressive. Importantly, integrinScore is cancer-dependently and closely associated with proliferation, stemness, tumor microenvironment, metastasis, and immune signatures. IntegrinScore also predicts patient’s response to immunotherapy. By mining drug databases, we unravel an array of compounds that may serve as integrin signaling modulators. Finally, we build a user-friendly database to facilitate researchers to explore integrin-related knowledge. Collectively, we provide a comprehensive characterization of integrins across cancers, which offers gene- and cancer-specific rationales for developing integrin-targeted therapy.

## INTRODUCTION

Integrins are a family of heterodimeric transmembrane glycoprotein receptors consisting of α and β subunits. In humans, 26 integrins are formed by sophisticated combinations of 18 α and 8 β subunits (Fig. 1A). Through binding to adjacent ligands, integrins act as adhesion receptors and bidirectionally transmit biochemical signals across the plasma membrane, thus enabling cells to rapid response to intracellular and extracellular cues. Integrins can adhere to nearly all extracellular matrix (ECM) components, rendering them an ability to remodel the extracellular environment. Depending on different types of ligands, integrins can be classified generally into four categories (Fig. 1A) [1, 2]. Especially, the same ligand can be bound by different categories of integrins, and the same integrin can also recognize and bind to multiple distinct ligands, creating an intricate network of integrin signalosome. Biologically, integrin signalosome represents a dynamic machinery responsible for various cellular “outside-in” and “inside-out” signal transmission and thus regulates a spectrum of important biological processes such as cell growth, survival, and migration [3]. Notably, dysregulation of integrin signalosome is tightly associated with the onset of many human diseases, including cancer [3].

**Figure 1.**
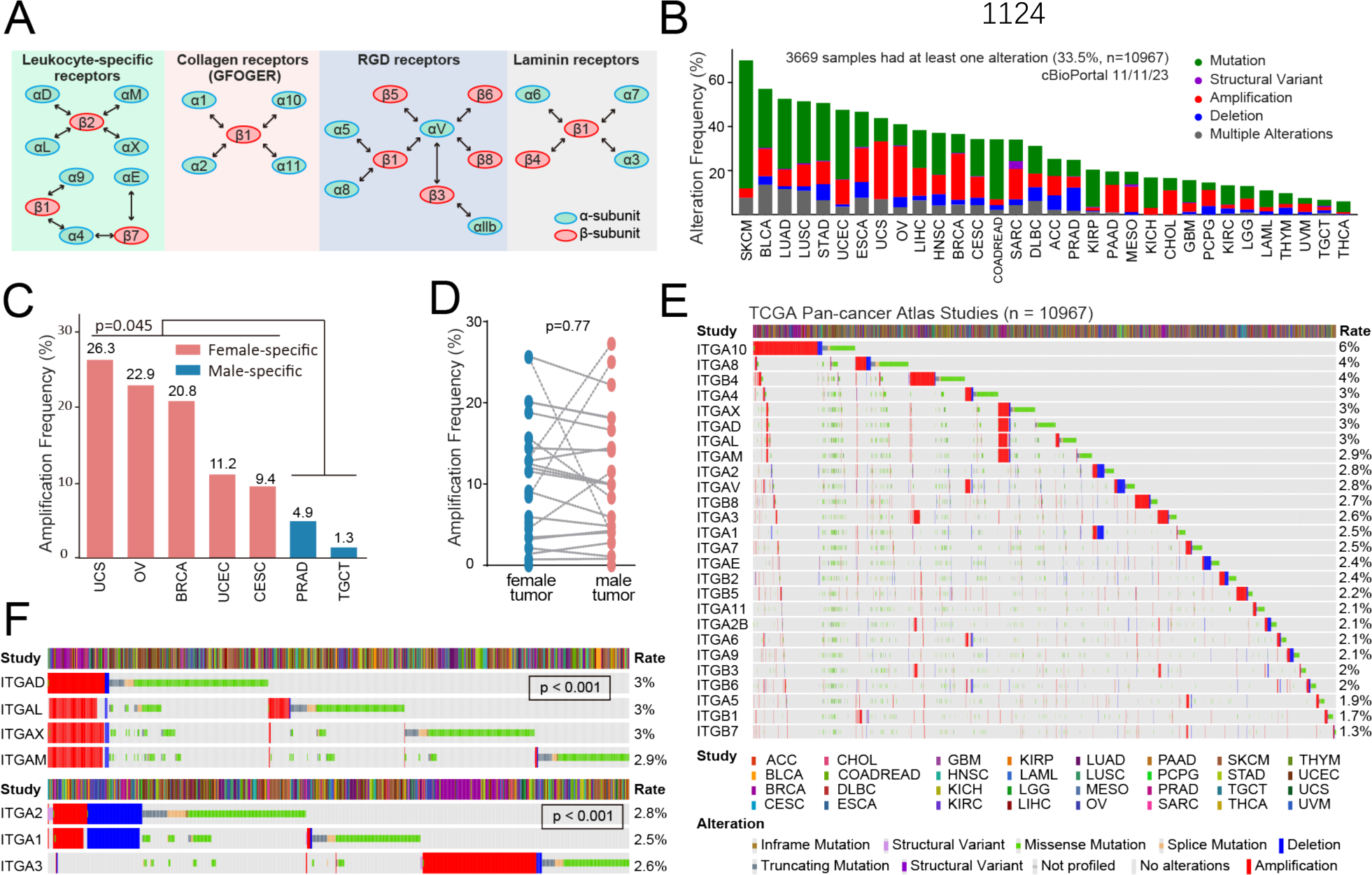
Integrin represents a frequently altered pathway at pan-cancer level. **A.** All 24 distinct integrin heterodimers formed by 26 integrin genes are classified into four groups according to cognate ligand. Bidirectional arrows are intended to demonstrate which integrin genes can form heterodimers. **B.** Cumulative alteration frequencies of genes encoding integrins in 32 human cancers. Shown are bar plots illustrating the cumulative alteration frequencies of all 26 integrin genes combined. **C.** Female-specific tumors have higher amplification frequencies than male-specific tumors. Student’s t-test is used for comparison. **D.** Comparison of amplification frequencies between female and male cancers. Student’s t-test is used for comparison. Female-specific (BRCA, CESC, OV, UCEC and UCS) and male-specific (PRAD and TGCT) tumors are not included. **E.** Alteration frequencies of 26 integrin genes in 32 human cancers. Shown are heatmaps illustrating distinct alteration frequencies of all 26 integrin genes individually. **F.** Co-occurrence and mutually exclusion pattern of seven integrin genes. Data shown in the right panel are alteration frequencies. P values are derived from Fisher’s exact test.

Evidence has implicated integrins in almost every step of cancer development and progression (including metastasis and drug resistance) [1, 4–6]. For example, comparative genomic analysis of tumors and matched normal tissues indicated that mutations of ITGA7 that cause protein truncations are associated with tumor initiation in prostate, brain and liver tissues [7]. In both hematological and solid cancers, studies have established integrins as key players in regulating proliferation, migration, and invasiveness via interaction with multiple oncogenic pathways [5, 6, 8]. As such, integrins have emerged as valuable therapeutic targets for drug development. Recently, we have provided our perspective on how integrin signalosome modulates cancer stemness, and summarized current clinical trials evaluating integrin-targeting drugs in solid tumors [4]. More broadly, dozens of integrin-targeted drugs are in the investigation in more than 130 clinical trials [1, 4].

Immunotherapy has revolutionized cancer treatment, evidenced by the discovery and successful clinical implementation of immune-checkpoint blockades (ICBs) targeting CTLA4, PD-1 and PD- L1 [9]. However, the majority of patients do not respond beneficially to ICB due to a number of mechanisms (such as the immune escape of cancer cells, inactivation of antitumor immune cells, and tumor microenvironment (TME) remodeling) that block cytotoxicity of T cells. Due to the intrinsic properties of modulating cell–cell and cell–ECM interaction, cancer-related integrins are reported to be tightly associated with immunosuppressive events listed above [10–12], providing a rationale for combining integrin inhibitors with ICB to achieve a better antitumor efficacy. In support, integrin αvβ6 can deactivate T cells and drive immune evasion and thus inhibition of αvβ6 in combination with ICB significantly improves patients’ responses to immunotherapy in colorectal [13] and breast cancers [14].

The tumor is an extremely heterogeneous tissue, with different genes and/or pathways exerting functions in a cancer-dependent manner. Therefore, understanding integrin-mediated roles and interactomes in distinct tumor types are prerequisite to devising better antitumor strategies. Despite recent advances of characterizing individual integrin’s role in cancer biology, a comprehensive and tumor type specific investigation of the whole 26 integrin genes is still lacking. Using the Cancer Genome Atlas (TCGA) database, which has profiled the genomic, transcriptomic and epigenomic landscape of nearly 10,000 primary tumors across 32 cancer types (Table S1) [15], we systemically analyze and compare the genomic and transcriptomic profiles of integrin pathway as a whole, as well as integrin interactomes, across distinct cancers. We also identify the molecular mechanisms underpinning the dysregulation of integrins. We report that integrin misexpression, caused by a combinatorial effect of genomic alteration, DNA methylation- and microRNA (miRNA)-mediated epigenetic regulation, and transcription factor (TF)-mediated transcription, and likely other fashions, plays generally an oncogenic role in most of the cancers. The expression profile of 26 integrin genes consistently identifies a subgroup of aggressive tumors featured by a high level of proliferation and stemness. An integrinScore, a weighted signature considering the dysregulation and prognostic values of integrin genes, is developed to facilitate our findings toward a potential clinical usage by quantifying the integrin pathway pattern in individual tumors. Besides a close correlation of integrinScore to patients’ outcomes and previously reported molecular subtypes in many cancers, integrinScore can be used to predict responses to ICBs. By mining the Connective Map (CMap) and the Genomics of Drug Sensitivity in Cancer (GDSC) drug sensitivity databases, we discover a number of potential drugs that might interfere integrin proteins for suppression or activation. Finally, we build a user- friendly database PIExplorer (http://computationalbiology.cn/PIExplorer), a valuable resource for the biomedical community to browse, search and download data of interest.

## RESULTS

### 1 Genomic alteration landscape of integrin genes in human cancers

To depict the pan-cancer genomic aberrations of genes encoding human integrins (hereafter referred to as integrin genes), we surveyed the alterations of all 26 integrin genes (Fig. 1A) across 32 cancer types via cBioPortal [16]. Aggregately, the integrin family as a group of cell adhesion molecules represented a frequently-altered pathway in 32 cancers. On average, 33.5% of the tumors contained at least one alteration of one integrin gene, with SKCM (70.0%) and THCA (6.0%) being the highest and lowest mutated cancer types, respectively (Fig. 1B, Table S2). Examination of details in distinct alteration forms across cancers revealed several interesting patterns. ***First***, the mutation occurred most often, followed by amplification, in the majority of cancers especially in SKCM (58.1%; Fig. 1B, Table S2), consistent with a high mutation burden for SKCM among human cancers [17]. ***Second***, amplification was observed more frequently in five female-specific cancers (UCS, OV, BRCA, UCEC and CESC) than in two male-specific cancers (PRAD and TCGT; Fig. 1C), while no significant difference was found between these two groups for mutation (t-test: p=0.089) and deletion (t-test, p=0.58). However, when we compared the amplification rate in the remaining non-sex-biased 25 cancers stratified by patient’s sex, we failed to observed a noticeable difference (Fig. 1D). These data strongly indicated that amplification in integrin genes may be associated with sex-hormone sensitive cancer types. ***Third***, deletion occurred predominately in PRAD (10.5%), STAD (7.3%) and ESCA (7.1%), indicating presumably tumor suppressive role of and loss-of-expression of integrin genes in these cancer types (also see below). ***Fourth***, despite a high alteration rate in integrin genes as a whole, most individual integrin genes were mutated at low frequency, with an average alteration rate of 3% (Fig. 1E). Interestingly, we found that several integrin genes were co-mutated. For example, ITGAD, ITGAL, ITGAX and ITGAM were all located at Chr16p11.2 and thus tended to be co-ampliated (p<0.001; Fig. 1F). Reports have been associated the copy number variants (CNVs) in Chr16p11 region with neurodevelopmental disorders and invasive BRCA [18]. Similarly, due to distinct genomic locations, ITGA1 and ITGA2 both resided at Chr5q11.2 were co-altered (p<0.001) but mutually exclusive to ITGA3, which is located at Chr17q21.33 (Fig. 1F). Integrin proteins function through heterodimers and are generally grouped into four categories (Fig. 1A) according to their ligand types [1, 4]. ITGAD, ITGAL, ITGAX and ITGAM all dimerize with ITGAB2 to form leukocyte-specific receptors. Therefore, the co-amplification of these four α integrins potentially indicated a functional similarity and/or redundancy among them, as well as biological importance, in modulating tumor immunity. Moreover, although all coupled with ITGB1, ITGA3 functions as a laminin receptor, whereas ITGA1 and ITGA2 function as collagen receptors. A mutually exclusive mutation pattern between ITGA3 and ITGA1/2 suggested potentially a differential requirement of them in cancer development. ***Fifth***, a detailed comparison of the mutational landscape of individual integrin genes across 32 cancers revealed that they altered in both a tumor- and gene-specific manner (Fig. S1A). For example, in terms of mutation rate, ITGA10 (6%) and ITGB7 (1.3%) are the highest and lowest altered ones, respectively (Fig. 1E). In terms of alteration types, ITGA10 had high amplification rates in LIHC (9.7%), BRCA (9.2%) and BLCA (9.0%), whereas it predominately mutated in SKCM (7.8%). In terms of cancer types, ITGA10 and ITGA5 represented the most frequently amplified genes in CHOL, while the other integrin genes were rarely amplified (Fig. S1A). Moreover, integrin genes are frequently, but rarely, depleted in PRAD and CHOL, respectively. In total, our data indicate that integrin genes collectively represent a frequently mutated pathway in human cancers. Importantly, we observed an overt heterogeneity in the alteration landscape of integrin family at multiple layers, suggesting diversity in mechanisms of action of the integrins underpinning cancer evolution.

Classification of pan-cancer patients with or without genomic alterations in integrin genes unraveled that the altered group displayed shorter overall survival (OS) and progression-free survival (PFS) time than the non-altered group (Fig. S1B and S1C), indicating a general pro- tumorigenic role for the gnomically altered integrin family. Particularly, when we “zoomed in” on individual genes, we found that: 1) alteration of most individual integrin genes was not prognostic across 32 cancers (only 3.0% for OS and 3.1% for PFS; Fig. S1D and S1E), which could be attributed to limit alteration events in distinct cancer types; 2) prognostic integrin genes based on genomic alterations displayed cancer type dependency, as many tended to be consistently pro-tumorigenic in HNSC but tumor-suppressive in UCEC (Fig. S1D and S1E). Collectively, the prognostic power of integrin genes based solely on genomic alterations was inadequate to stratify cancer cohorts clearly.

### 2 Expression dysregulation of integrin genes across cancers

We next examined the global misexpression of the integrin family by interrogating the transcriptomes of 15 TCGA cancer types, which have at least 10 matched tumor and normal samples for each cancer type. Results showed that, globally, all integrin genes were deregulated in the 15 cancers, with each gene misexpressed in at least 4-12 cancers (Fig. 2A, Table S3). Notably, both cancer-type-specific and gene-specific dysregulation were seen. LUSC had the most deregulated integrin genes (21 genes), followed by KIRP (18 genes) and KICH/HNSC (17 genes) (Fig. 2A, bottom). Particularly, HNSC, THCA and KIRC represented cancers with preferentially upregulated integrin genes, whereas PRAD, UCEC and KICH showed an opposite pattern (Fig. 2A, bottom). At the individual gene level, ITGA8 and ITGA9 were down-regulated in 11 out of 15 cancers, whereas ITGAX, ITGB4 and ITGA6 were more frequently up-regulated in tumors than adjacent normal tissues (Fig. 2A, right). These results together highlighted a heterogeneity in the dysregulation of integrin genes across human cancers, reflecting distinct roles played by integrins in different pathological contexts. Of note, consistent with their co-amplification profile of ITGAD, ITGAL, ITGAX and ITGAM (Fig. 1E), we observed similar dysregulation patterns for them in cancers (belonging to one sub-cluster, Fig. 2A), validating our analytic pipeline.

**Figure 2.**
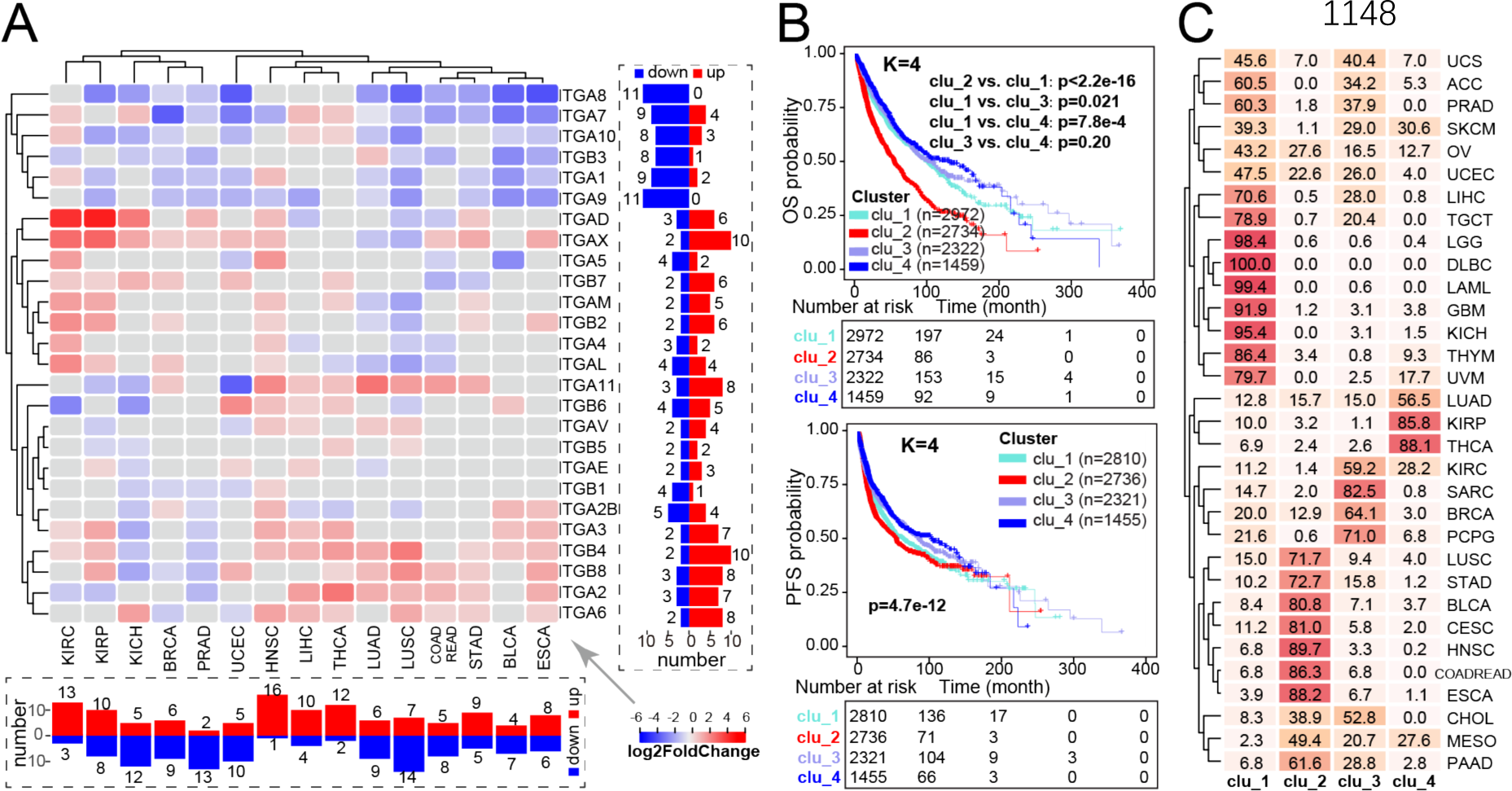
Integrin genes dysregulate in tumors compared with adjacent normal tissues and their general expression patterns correlate with patient’s outcome. **A.** Heatmap showing differential expression of integrin genes between tumors and adjacent normal tissues. The right and bottom panel summary the number of significantly changed integrins at gene and tumor level, respectively. Grey rectangles denote unsignificant results at the cutoff of absolute of FC > 1 and FDR < 0.05. Unsupervised clustering method is used. **B.** Four integrin expression patterns correlate with patient’s OS (the top two panels) and PFS (the bottom two panels) time. **C.** Sample distribution in the four clusters. Data and color shown in boxes are the percentage and intensity of samples classified in corresponding clusters, respectively.

Cancer-specific dysregulation of integrin genes indicated potentially a population-stratification ability according to their expression profile. Using unsupervised consensus clustering, we grouped all 32 pan-cancer samples according to the transcriptome of 26 integrin genes into four subclusters (k=4; namely clu_1 to 4; Table S4), with clu_2 showing the shortest OS and PFS survival followed by clu_1 (Fig. 2B). Especially, to ensure k=4 was the best stratification (Fig. S2A), we also tested k=3 and k=5 and observed similar results, with clu_2 also being the clinically aggressive cluster (Fig. S2B-S2G). Importantly, regardless of the cutoff of subcluster number (k=3, 4, or 5), the dysregulation of integrin genes identified, convergently, the same subgroup of tumors with worse outcomes (Fig. S2D and S2G). Examining of cancer types belonging to the four subclusters, we found that DLBC (100%), LAML (99.4%) and LGG (98.4%) were mainly resided in clu_1; whereas HNSC (89.7%), ESCA (88.2%) and COADREAD (86.3%) predominately belonged to clu_2 (Fig. 2C), in line with the cancer-dependent transcriptome of integrin genes. As such, different subclusters were dominated by distinct integrin genes (Fig. S2H). For example, ITGB4, ITGB6, ITGA2 and ITGA6 predominantly expressed in clu_2; while ITGA9, ITGB3 and ITGA3 expressed mainly in clu_4. Considering the fact that ITGB4, ITGA2 and ITGA6 were frequently up-regulated in cancer vs. normal tissues (Fig. 2A), our results suggested potential pro-tumorigenic roles for them. Indeed, oncogenic roles have been reported for these three integrins in different cancer types [4].

### 3 Mechanisms underpinning transcriptional integrin dysregulation

Genomic (Fig. 1) and transcriptional (Fig. 2) aberrations underscore the significance of the integrin family in human cancers. Then, a question that arises is what mechanisms drive their expression dysregulation. We considered both genetic and epigenetic mechanisms. ***First of all***, genomic alterations are major drivers of cancer initiation and progression [19]. CNVs are frequently linked to altered gene expression [20]. Indeed, amplification and deletion generally correlated with gain and loss of mRNA expression, respectively (Fig. 3A). It is noteworthy that the mutational burden for individual integrin genes was quite low in pan-cancer samples (Fig. 1E), suggesting that genomic alteration could only explain in small part the mis-expression of them and thus the existence of other regulatory mechanisms.

**Figure 3.**
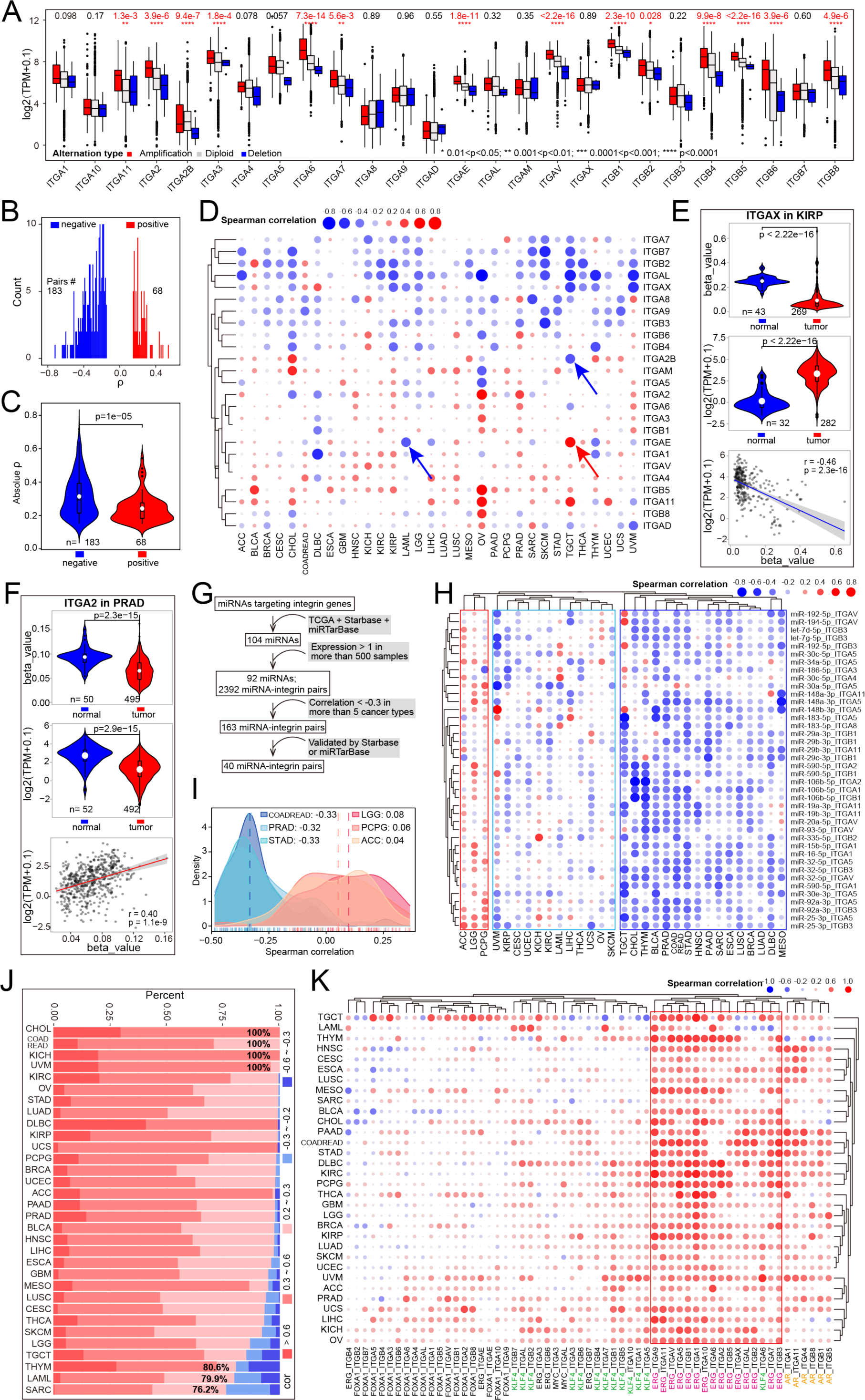
Mechanisms underlying integrin dysregulation in tumor. **A.** Boxplots showing integrin expression in tumor samples with amplification, diploid and deletion alteration, with p values shown on the top. Within the plots, the center lines represent median values, box edges are 75th and 25th percentiles, and dots denote the outliers. **B.** Bar plots showing the number of integrin-probe pairs with varied P values. 183 and 68 significantly negatively and positively correlated pairs are shown, respectively. **C.** Violin plots showing the absolute p values in negatively and positively correlated pairs. The Wilcoxon test is used. Within the violin plots, the center white dots represent mean values, box edges are 75th and 25th percentiles, and dots denote the outliers, so as to the **E** and **F** plots. **D.** Spearman correlation of DNA methylation in promoter with expression of integrin genes. Blue and red arrows denote three examples for distinct correlation between integrin-probe pairs. **E and F.** DNA methylation partially contribute to ITGAX (**E**) and ITGA2 (**F**) dysregulation in tumors. Top and middle: violin plots showing DNA methylation in promoter and expression of integrin gene in tumors and normal tissues, respectively; bottom: dot plots showing Spearman correlation between DNA methylation in promoter and expression of integrin genes. **G.** Workflow for correlation analysis between miRNA and integrin genes. **H.** Spearman correlation of expression of miRNAs and integrin genes. Three correlation patterns (negative-, mixed- and positive-relationship) are identified by unsupervised clustering method. **I.** Density plots showing correlation coefficient between miRNAs and integrin genes in extreme cancers. Three positive-correlation dominant cancers (LGG, PCPG and ACC) and three negative-correlation dominant cancers (COADREAD, PRAD and STAD) are shown. **J.** Bar plots showing cumulative percentage of significant correlated transcription factor (TF)-integrin pairs with distinct range of correlation coefficient. The cutoff of Spearman ρ ≥ 0.2 and P < 0.05 is considered significant. The cancers are ordered by the percentage of positively correlated pairs. Several extreme cancers are labeled with the percentage values. **K.** Spearman correlation of expression of TFs and integrin genes. The relationships between ERG and integrin genes are overwhelmingly positive and boxed by red square. For **D**, **H** and **K**, positive and negative correlations are colored in red and blue, respectively. The size of dots indicates the absolute correlation coefficient.

***Next***, we examined the impact of DNA methylation on gene expression. By focusing on the DNA methylation probes within promoter regions of integrin genes, we observed a significant inverse correlation for the majority of integrin-probe pairs in pan-cancer, with a cutoff of absolute Spearman’s correlation (ρ) ≥ 0.15 (Fig. 3B), suggesting, as expected, that DNA methylation (measured as probe β-value) suppressed gene expression. Moreover, we also noticed that the absolute ρ values for negative integrin-probe pairs were larger than that of positive pairs (Fig. 3C), indicating a severity in inhibiting, rather than enhancing, integrin gene expression by DNA methylation. A positive correlation between DNA methylation and elevated gene expression was also reported [21]. When we profiled the relationship between DNA methylation and gene expression for individual integrin genes in individual cancer types, we observed gene- and also cancer-dependent patterns (Fig. 3D, Table S5). For example, DNA methylation of ITGA2B and ITGAE negatively and positively correlated with their expression in TGCT, respectively. Also, at gene level, an opposite relationship was found for the same ITGAE gene in different cancer types (i.e., TGCT and LAML; Fig. S3A). Integrative analysis combining transcriptome and DNA methylome revealed that DNA methylation may explain the dysregulation of some but not all integrin genes. ITGAX had a lower promoter methylation level in, and thus displayed overexpression in, tumor vs. normal tissues in KRIP (Fig. 3E). For ITGA2, a positive correlation between its expression and methylation was observed, in line with both lower expression and methylation level in tumor vs., normal tissues in PRAD (Fig. 3F). Similarly, inconsistency between expression and methylation level for ITGA11 in LIHC (Fig. S3B) and ITGB7 in THCA (Fig. S3C) was also noticed. Both elevated expression and methylation were observed for ITGB7 in THCA, indicating that methylation may enhance expression, but instead, an inverse correlation between them was found (Fig. S3C). Collectively, our data illustrated a cancer- dependent effect of DNA methylation on regulating integrin gene expression, and the disagreement between integrin misexpression and mis-methylation highlighted the involvement of other regulatory layers.

***Third***, we investigated the miRNA-mediated repression of integrin genes, as dysregulated miRNA profile contributes to tumor development via reshaping cancer transcriptomes [22–24]. To comprehensively profile the miRNA regulome for integrin genes, we first constructed a strict analysis workflow (Fig. 3G) to narrow down to the final 40 miRNA-integrin pairs. Results indicated that the majority of the miRNA-integrin pairs in TCGA cohorts had a negative relationship with a median correlation of -0.16 (Fig. S3D, Table S6). Such association between miRNA-integrin pairs was also validated in other independent dataset (Fig. S3E). Utilizing unsupervised clustering, we obtained three pan-cancer subgroups marked by negative-, mixed- and positive-relationship among miRNA-integrin pairs (Fig. 3H). By a cutoff of absolute Spearman ρ ≥ 0.3, COADREAD had the highest number (n=28) of negatively correlated miRNA-integrin pairs, followed by PRAD (n=25) and STAD (n=24); whereas LGG, ACC, and PCPG represented three exceptional cancers which predominantly harbored positively correlated miRNA-integrin pairs (Fig. 3H, 3I and S3F). Interestingly, PRAD and BLCA, both originated from the genitourinary system, exhibited similar miRNA-integrin correlation patterns (Fig. 3H). However, COADREAD and LGG showed nearly opposite correlation patterns (Fig. 3H), as the same set of miRNA-integrin pairs produced inverse outcomes in terms of modulating gene expression (Fig. S3G and S3H). This cancer type specificity may be reflected by the intrinsic abnormities in miRNA biogenesis and maturation processes in distinct cancer types. For example, it has been reported that malfunction of the posttranscriptional maturation of miRNAome caused by aberrant nuclear localization of DICER could lead to widespread dysregulation of miRNAs expression, as well as a consequence miRNAs do not efficiently target their targets, in glioblastoma [25]. In support, we observed that, for certain miRNAs, a substantial proportion of targeting genes (83.3% for miR-32-5p and 64.0% for miR-148a-3p) were positively rather than negatively correlated with their expression in LGG (Fig. S3I). As such, this global miRNAome abnormity may contribute to the twisted miRNA-integrin correlations seen in LGG vs. other negative-relationship cancer types (Fig. 3H and 3I).

***Last***, we investigated a transcription factor (TF)-based mechanism, as oncogenic TFs drive nearly all cancer-related biological processes by controlling transcription [26, 27]. In a human TF database, 495 human TFs were annotated with potential targets [28]. By retrieving all TFs that potentially target integrin genes, we chose the top ten TFs with the most integrin targets for further analysis (Fig. S3J). By a cutoff of absolute Spearman ρ ≥ 0.2, we found that positively correlated TF-integrin pairs constituted the vast majority of significant ones (Fig. 3J, Table S7), suggesting an overall transcriptional promotion on integrin genes by these TFs regardless of cancer types. Many TFs are oncogenic drivers. For example, MYC is a notorious oncoprotein overexpressed in the majority of human cancers [29], and AR (androgen receptor) and FOXA1 are pioneer factors in PRAD [30]. Focusing on five well-studied oncogenic TFs (i.e., AR, FOXA1, ERG, KLK4, and MYC), we profiled their potential regulation on integrin genes, and found a positive correlation between the expression of most TF-integrin pairs (Fig. 3K, Table S7). Relatively, although FOXA1 and MYC had less significant correlation with integrin genes, ERG showed an overwhelmingly positive correlation with them in pan-cancers, pointing ERG as an essential upstream TF of integrins. AR is expressed in multiple tissues, and it positively correlated with six integrin genes in PRAD and also with other integrin genes in other cancer types (e.g., HNSC, CESC), indicating a weak cancer type specificity. KLF4 is a renowned stemness factor [31]. The moderate positive correlation between KLF4 and integrin genes in pan-cancers implied that KLF4 may impact cellular stemness by, at least partially, regulating integrin genes. In support, evidence has implicated integrins in regulating cancer stemness [4]. We also observed several negatively correlated TF-integrin pairs, such as FOXA1-ITGB4 in PRAD and FOXA1-ITGA3/ITGA5 in HNSC (Fig. 3K). Consistent with their negative correlation of expression, an opposite expression pattern for these FOXA1-integrin pairs in both PRAD and HNSC was found (Fig. S3K). To further solidify our findings, we also interrogated another two independent datasets, GSE157548 for PRAD and the Chinese Glioma Genome Atlas (CGGA) [32] for LGG, for expression correlation between TFs and integrin genes. Similar to that was found in TCGA cohorts, these TFs were expected to play strong regulatory roles in modulating their expression (Fig. S3L and S3M). Our data together illustrated a notion that there is an array of TFs, but unlikely a single TF, to cooperatively determine an integrin profile in each cancer type.

In summary, we concluded that an intricate network of mechanisms (including genomic alteration, DNA methylation, miRNA, dominant oncogenic TFs, and perhaps others) is responsible for the transcriptional dysregulation of integrin genes in pan-cancer and that such a network operates likely in a cancer- and gene-dependent manner. These mechanisms represent different regulatory layers of integrin gene regulation. Conceivably, cross-talk between them is expected. For example, after integrating the associations among miRNA, TF and integrin genes, we depicted a reciprocal and cross-regulatory map (Fig. S3N, left). Specifically, both miR-29b-3p and MYC could regulate ITGB1, and in turn MYC may also modulate miR-29b-3p (Fig. S3N, right). To experimentally validate the impact of different mechanisms (i.e., miRNA, DNA methylation, TF) on integrin gene expression, we utilized AR^+^ LNCaP as a model representing PRAD. In our pan-cancer analysis, miR-92a-3p was identified as a node miRNA regulating integrin family (Fig. 3H). Thus, 5’-Aza (a DNA methylation inhibitor), miR-92a-3p overexpression, and Enzalutamide (an AR inhibitor) were used to disrupt the global DNA methylation, miRNA expression, and a prominent TF’s activity, respectively. As for a readout, we chose the indicated 12 integrin genes whose expression was really detectable in LNCaP cells. Results showed that expression of many of the integrin genes has altered after 5’-Aza (Fig. S3O), miR-92a-3p overexpression (Fig. S3P), or Enzalutamide treatment (Fig. S3Q), demonstrating that modulation of these regulatory mechanisms could result in dysregulation of integrin family in cancer cells.

### 4 Integrin expression patterns classify pan-cancers into four clusters with distinct clinical features

To molecularly characterize our four integrin subclusters (Fig. 2B), we utilized two recently reported pan-cancer classifications. Thorsson et al. have integrated major immunogenomics methods to classify >10,000 TCGA samples into six immune subtypes (namely C1 to C6), of which C1 and C2 were marked by enhanced proliferation [33]. Here, we found that more than half of the tumors originally grouped in C1 (53.8%) and C2 (52.5%) were classified as clu_2 in our study (Fig. 4A). Notably, GSVA against proliferation-related signatures showed that tumors in clu_2 possessed the highest scores vs. other subclusters (Fig. 4B), highlighting a proliferative nature. In our four integrin subclusters, clu_2 and clu_1 represented the top two deadly subclusters according to the survival analysis (Fig. 2B). Stemness is a hallmark of cancer and parallels with cancer progression [34]. Recently, a pan-cancer stemness feature correlated with oncogenic dedifferentiation and metastasis has been identified via machine learning on large- scale molecular data [35]. Interestingly, we found that clu_1 and clu_2 displayed a higher stemness index among the four subclusters (Fig. 4C), in line with their worse clinical outcome (Fig. 2B). Collectively, the transcriptome of integrin family could stratify pan-cancer patients into four subclusters with distinct clinical and molecular features. Particularly, tumors in clu_2 exhibit higher proliferation and stemness and thus have worse outcomes.

**Figure 4.**
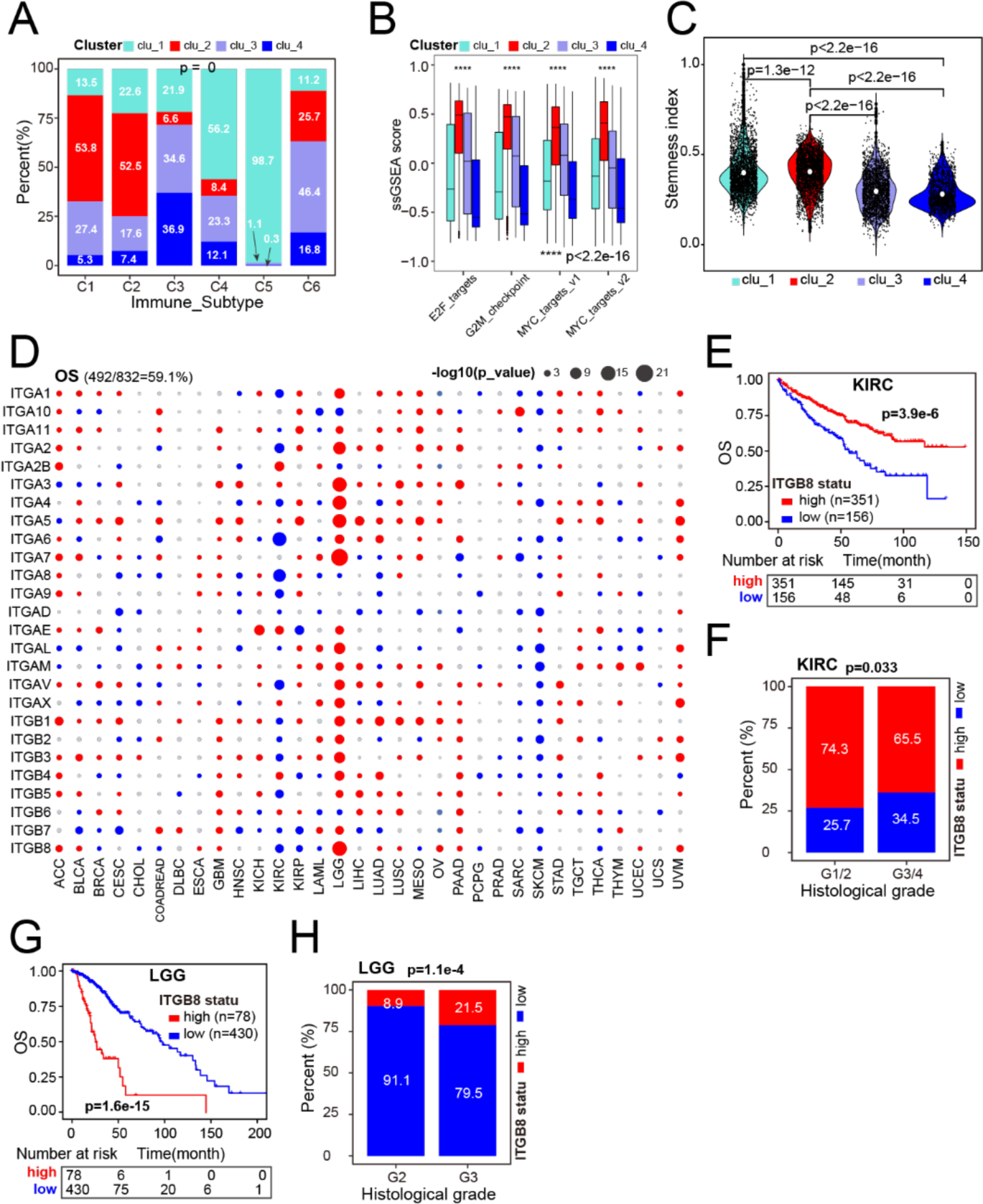
General expression patterns of integrin genes correlate with cancer hallmarks and patient’s survival. **A.** Distribution of six previously reported immune subtypes in four clusters, with p value labeled on top. **B.** Boxplot quantification of the activity of proliferation related signatures among four clusters. Within the plots, the center lines represent median values, box edges are 75th and 25th percentiles, and dots denote the outliers. **C.** Violin plots showing the stemness index among four clusters. Within the plots, the center white dots represent mean values. P values are calculated by the Wilcoxon test. **D.** Summary of overall survival (OS) time analysis results of integrin genes across cancers. Red, blue and grey dots indicate worse, better and unsignificant results, respectively. The size of dots indicates -log2(P_values). Data in the parentheses (top left) denotes the rate of dysregulation associated with OS. **E** and **G.** Kaplan-Meier plots illustrating ITGA8 as a favorable and an unfavorable gene associated with patient OS in KIRC and LGG, respectively. **F** and **H.** Comparison of the percentage of distinct histological grades between ITGA8 high- and low-expression group in KIRC and LGG, respectively. For **A**, **F** and **H**, chi-square test is used.

Being prognostic as a whole, we next assessed the prognostic values of 26 individual integrin genes in patient’s outcome. We found that 59.1% and 58.1% of dysregulated integrin genes frequently correlated with patients’ OS (Fig. 4D) and PFS (Fig. S4A) in distinct cancer types (Table S8), respectively. Globally, integrin genes with higher expression generally acted as risk factors across cancers (Fig. 4D), especially in LGG, ACC, and UVM. In contrast, higher expression of the majority of integrin genes uniformly correlated with a longer OS/PFS in KIRC, SARC, and SKCM, indicating them as protectors in specific cancer types. Interestingly, for the same integrin genes, a heterogeneous correlation with patients’ OS/PFS was also observed. For example, ITGB8 may play a tumor suppressive role in KIRC, as patients whose tumors had higher ITGB8 expression displayed longer OS time and lower tumor stage (Fig. 4E and 4F). But, conversely, ITGB8 might play an oncogenic role in LGG (Fig. 4G and 4H). ITGA5 was associated with a worse patient’s outcome in most of cancer types (19/32), but its higher expression correlated with a good prognosis in ACC, indicating a protective role (Fig. S4B). Similar pattern was also observed for ITGB7 (Fig. S4C). Interestingly, for cancers originated from the same tissue (like kidney cancers KICH, KIRC, and KIRP), the same integrin genes may play opposing roles in different contexts. For example, ITGAE was proposed to be oncogenic in KICH and KIRC, but tumor suppressive in KIRP (Fig. S4D). In aggregate, although a large number of integrin genes were prognostic, we did observe a gene- and cancer-specific pattern, highlighting a need to develop a formula that considers and simplifies such heterogeneity before translating the prognostic values of integrins into the clinic.

### 5 integrinScore molecularly separates aggressive primary tumors from indolent ones

The widespread dysregulation of integrin genes possessed prognostic values either at the family (Fig. 2B) or individual gene (Fig. 4D) level across cancers. However, the directional heterogeneity in both misexpression (up- or down-regulated) and prognostic values (risk factor or protector) across cancers hindered us from accessing and integrating all integrin genes in individual patients for potential clinic use by traditional methods like GSVA [36] or Z-normalization [37]. Here, we adopted a method reported previously by us [38] to construct integrinScore, a weighted scoring model that considered both the prognostic value (risk and protective gene as positively and negatively weighted, respectively) and normalized expression value (up- or down-regulated in a given tumor type) of all integrin genes in a tumor sample (see Methods). Survival analysis revealed that integrinScore well-separated patients into two groups, with the integrinScore high group having a shorter survival than the low group, as expected (Fig. S5A). Using the same model, we extended our analysis to several other datasets containing 6 cancer types and yielded consistent results (Fig. S5B). Thus, integrinScore can be used as a biomarker to uniformly stratify a cohort of tumors with differential aggressiveness.

To strengthen our findings, we next performed association studies between integrinScore and a repertoire of previously reported clinical and/or molecular cancer features. Globally, integrinScore was significantly higher in tumors with high pathological t-stages (Fig. 5A), in line with a risk role of integrinScore in most of the cancers (Fig. S5A). Recently, Bagaev et al. categorized TCGA tumors into four TME subtypes (based on a set of 29 knowledge-based functional gene signatures): immune-enriched and fibrotic (IE/F), immune-enriched but nonfibrotic (IE), fibrotic (F) and immune-depleted (D) [39]. Examination of Thorsson’s six immune subtypes [33] and Bagaev’s four TME subtypes [39] with respect to integrinScore, we noticed several intriguing points. ***First***, integrinScore significantly fluctuated across both immune (15/27; Fig. 5B) and TME (22/24; Fig. 5C) subtypes in the majority of cancers, establishing a TME reshaping ability of integrins. ***Second***, the C1 subtype was reported to be more proliferative [33], but we observed that C1 tumors exhibited higher integrinScore only in a limited number of cancers (e.g. LIHC and LUAD), with some cancers in C1 (e.g., LUSC and STAD) displaying a lower integrinScore (Fig. 5B). In support, using two proliferation signatures, we found that integrinScore positively correlated with proliferation in LIHC and LUAD, but negatively correlated in LUSC and STAD (Fig. S5C), indicating a cancer-dependent relationship between integrinScore and proliferation. ***Third***, we found that among four TME subtypes at the pan- cancer level, integrinScore was discriminately associated with a representative IE-phenotype in some cancers, but with a F-phenotype in some other cancers (Fig. 5C). In the IE-phenotype cancers, such as ACC (Fig. S5D left), similar and lower integrinScore values were found for IE and IE/F subgroups. Combined stratification of ACC by integrinScore and IE-phenotype further identified an aggressive subgroup marked by integrinScore high and non-IE-phenotype (Fig. S5D right). In contrast, in F-phenotype cancers, took OV as an example, tumors in F and IE/F showed comparable high integrinScore values relative to D and IE (Fig. 5C, Fig. S5E left), and a combination of integrinScore high and F-phenotype further marked an aggressive tumor subgroup (Fig. S5E right). Notably, not all cancer subtypes can be separated by intergrinScore, as there was little difference in integrinScores for several cancers when classified by immune or TME features (Fig. 5B and 5C). For example, no change was found in integrinScores in UCS and PCPG subtypes.

**Figure 5.**
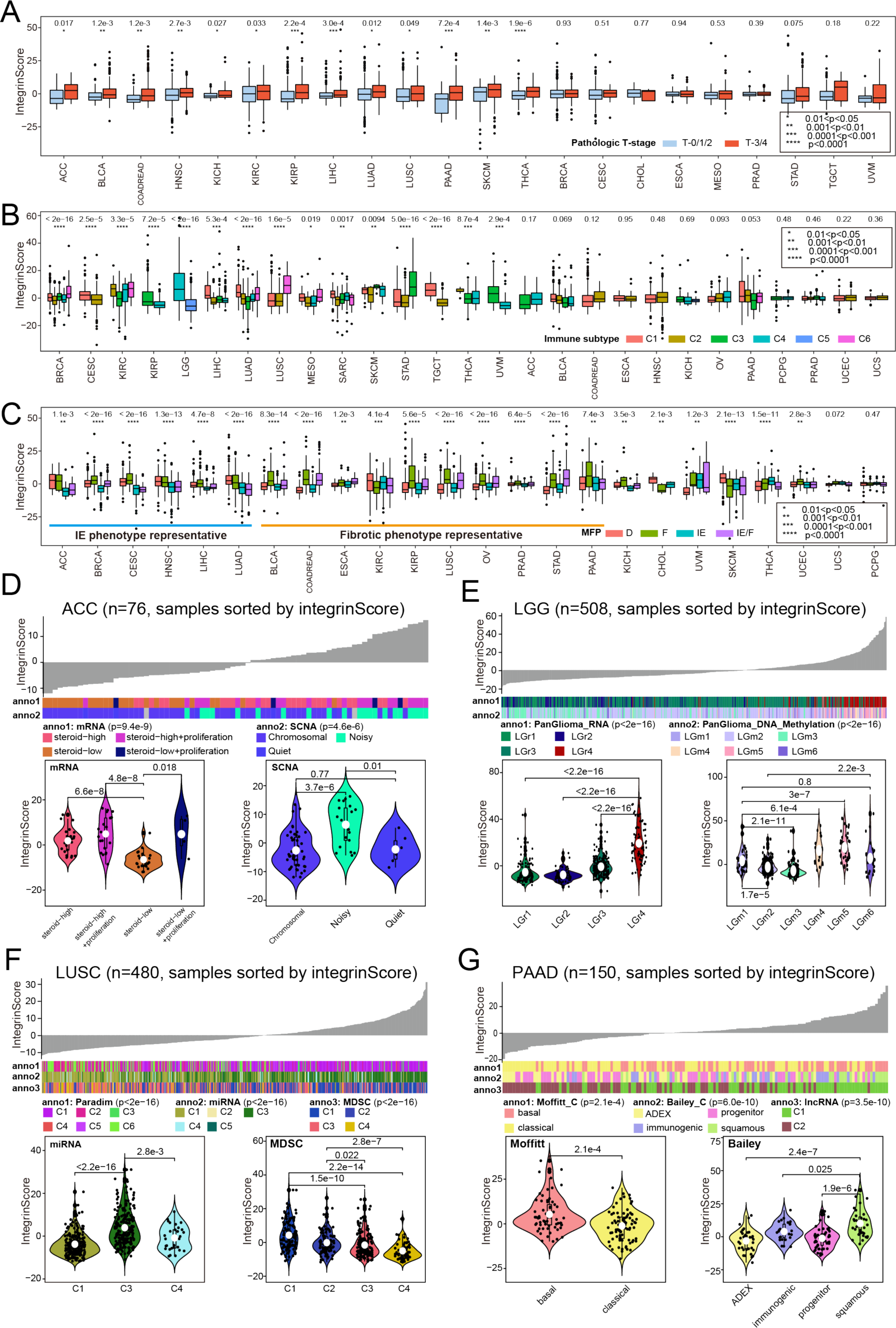
Molecular and clinical features associated with integrinScore in distinct cancers. **A.** Boxplots showing integrinScore values in higher (T-3/4) and lower (T-0/1/2) pathological T-stage across cancers. **B** and **C**. Boxplots showing integrinScore values among six immune subgroups (**B**) and four TME subgroups (**C**) across cancers. In **C**, IE and fibrotic phenotype representative cancers are labeled, respectively. **D-G**. Top: an overview of the association between known molecular subtypes and integrinScore in ACC (**D**), LGG (**E**), LUSC (**F**) and PAAD (**G**). Columns represent samples sorted by integrinScore from low to high. Bottom: violin plots showing integrinScore among distinct molecular subtypes. Within the boxplots and violin plots, the center lines represent median values, box edges are 75th and 25th percentiles, and white dots in violin plots denote mean values. P values are calculated by the Wilcoxon (for two groups) or Kruskal–Wallis (for more than two groups) methods.

To further illustrate the utility of integrinScore, we focused on individual cancer types. Generally, we observed a tight correlation between integrinScore and several known molecular subtypes for the majority of the TCGA cancers (Fig. 5D-5G and S5F-S5I). In ACC tumors, steroid hormones rendered a hostile environment to immune infiltrating cells and thus promoted tumor progression [40]. Consistently, we revealed that tumors with lower steroid levels tended to have lower integrinScore (Fig. 5D). It has been reported that a subgroup of ACC tumors (noisy group) characterized by a significantly higher number of chromosomal breaks due to frequent arm-level CNVs possessed worse prognosis [40]. Here we found that tumors in the noisy group had higher integrinScore (Fig. 5D). In LGG, Ceccarelli et al. have profiled more than 1100 diffuse gliomas into distinct molecular classifications by multidimensional omics [41]. They determined tumors in the PanGlioma_RNA cluster LGr4, and PanGlioma_DNA_methylation cluster LGm4/5/6 consisted of a large proportion of high-grade tumors with worse survival. Consistently, higher integrinScore values were seen in LGr4 and LGm4/5/6 tumors (Fig. 5E). Relevantly, classical (CL) and mesenchymal (ME) tumors, which exhibited worse outcomes, had higher integrinScore compared with neural (NE) and pro-neural (PN) ones (Fig. S5F, left). Combination of integrinScore with CL or ME subtype further stratified and thus identified a subgroup of aggressive tumors (Fig. S5F, right). Analysis of advanced brain malignancy GBM yielded similar results (Fig. S5G).

Based on ∼1000 variable pathway features, Campbell et al. have clustered >1300 squamous cancers (including LUSC) into 6 subclusters (PARADIGM C1 to C6), with C1 and C2 being enriched for active immune-related (i.e., IE-phenotype) and proliferation pathways, respectively [42]. LUSC was an F-phenotype representative cancer with F and IE/F subtypes showing higher integrinScore (Fig. S5H left). We noticed that tumors in PARADIGM C1 had higher integrinScore (Fig. S5H middle), so we speculated that PARADIGM C1 might contain more IE/F tumors. Projecting tumors in PARADIGM subclusters into four TME subtypes revealed that C1 had the highest proportion (29.2%) of IE/F tumors (Fig. S5H right), validating our hypothesis. We also found that tumors in proliferative C2 had lower integrinScores, in line with a negative correlation between proliferation and integrinScore in LUSC (Fig. S5H middle and S5C). IntegrinScore also showed a strong correlation with the miRNA cluster, with C3 having the highest value (Fig. 5F). Indeed, miRNA cluster C3 had more F and IE/F tumors (Fig. S5H right). Myeloid-derived suppressor cells (MDSCs) were a heterogeneous group of immune suppressive cells originating from the myeloid lineage, and the abundance of MDSC infiltration in TME correlated with worse outcomes and drug resistance [43, 44]. Based on 49 MDSC-related gene expressions, squamous cancers were clustered into four subclusters (MDSC C1 to C4), with C1 and C2 being the MDSC-inflamed subclusters associated with poor outcomes [42]. In our study, integrinScore of tumors decreased gradually from MDSC C1 to C4 (Fig. 5F), and the same trend was true for the proportion of F and IE/F tumors (Fig. S4H right), indicating that F-phenotype played an oncogenic role in LUSC.

A dense fibrotic stroma is the definitive feature of PAAD, which represent a barrier to captureing the intrinsic molecular features of epithelial cancer cells. In Moffitt’s study, a blind source separation method was utilized to classify PAAD tumors into basal and classical subtypes, of which basal PAAD subtype molecularly resembled basal breast tumors [45]. Consistently, basal PAAD tumors also had worse outcomes and higher integrinScore (Fig. 5G). Alternatively, Bailey et al. utilized an mRNA expression profile to define PAAD into four subtypes, with the squamous subtype exhibiting the worst outcome [46]. Here we found that squamous tumors had the highest integrinScore (Fig. 5G). Immunogenically, squamous tumors were characterized by the activation of CD4^+^ regulatory T cells and inhibition of antigen presenting cells, leading to immunosuppression in TME [46]. This phenotype was also reflected by a higher proportion of tumors in the D and F subtypes (Fig. S5I).

Besides aforementioned cancer types, we also extended our analysis to several other TCGA cancers, and found that integrinScore was also tightly correlated with multiple known subtypes defined by genomic (e.g., BRAF for THCA, SCNA for TGCT), transcriptomic (e.g., mRNA for KIRP and UCEC, miRNA for COADREAD and MESO, lncRNA for KIRC, PanGyn for OV and CESC, etc.), epigenetic (e.g., methylation for THCA and SKCM), and histologic (e.g., UCEC) features (Fig. S5J-S5W). Collectively, our data established integrinScore as a general marker capable of separating patients into clinically distinguishable subgroups. Importantly, the cancer type-specific but a consistent association of integrinScore with distinct previously reported molecular subtypes highlighted the functional diversity of integrins in tumor development and progression.

### 6 Molecular pathway characterization of pan-cancer integrinScore subclusters

Our findings that integrinScore uniformly identified a clinically aggressive cancer subcluster with distinct molecular features in a cancer type-specific manner prompted us to explore the biological pathways associated with the expression profile of integrin genes across cancers. We first identified the differentially expressed genes (DEGs) between integrinScore high vs. low groups and observed substantial variation in DEG number in different cancer types, ranging from 463 in CHOL to 8462 in PAAD (Fig. S6A). Certain cancer types had a high ratio of up- vs. down-regulated DEGs (e.g., 25.4 fold for STAD, 13.2 fold for KIRC, and 9.4 fold for ESCA); whereas an opposite pattern was also seen in HNSC (13.0 fold), CHOL (5.1 fold) and UCEC (4.3 fold), implying different roles of integrins in reprogramming cancer-specific transcriptomes. Based on the differential transcriptomes impacted by integrins, we then performed gene set enrichment analysis (GSEA) to identify key biological processes. Globally, cancer hallmarks were either positively or negatively enriched in integrinScore high (vs. low) groups in a cancer- dependent manner (Fig. 6A, Table S9). In most cancers, proliferation related pathways (e.g., E2F targets and G2M checkpoint) were positively enriched in integrinScore high subclusters (Fig. 6A). However, stemness related signatures such as MYC targets displayed both positive and negative enrichment in integrinScore high groups in a similar number of cancer types, indicating intrinsic biological difference in distinct cancer types. Notably, we noted that TME- related pathways (e.g., TGFβ signaling, epithelial mesenchymal transition (EMT) and angiogenesis), as well as immune signatures (e.g., TNFα signaling via NFkB, inflammatory response, and IL6-JAK-STAT3 signaling), were predominately positively, rather than negatively, enriched in integrinScore high groups across cancers.

**Figure 6.**
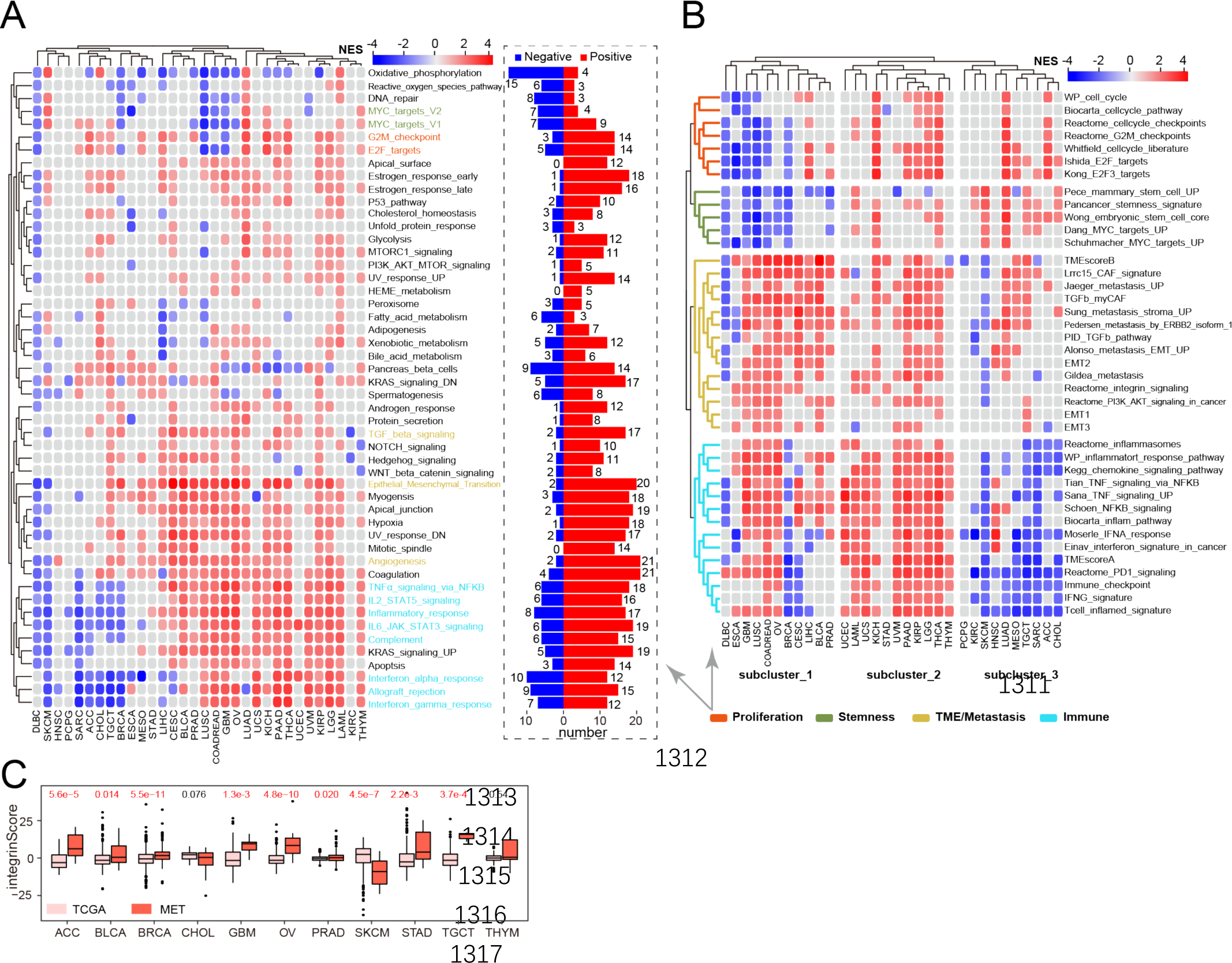
IntegrinScore correlates with cancer hallmarks across cancers. **A.** Left: heatmap showing NES, normalized enrichment score of 50 cancer hallmarks across cancers. Right: bar plots showing the number of cancer types in which each signature significantly enriches. **B.** Heatmap showing NES of curated signatures across cancers. Unsupervised clustering method is used for subgroup identification. **C.** Boxplots showing integrinScore in primary and metastatic tumors across cancers. P values are calculated by the Wilcoxon method. For **A** and **B**, red, blue and grey boxes denote positive, negative and unsignificant enrichment. Proliferation, stemness, TME/metastasis and immune related signatures are labeled with different colors.

To further solidify our findings, we performed GSEA against a curated sets of gene signatures (Table S10). Overall, a similar pattern of gene sets enrichment across cancers to that was shown in Fig. 6A was observed (Fig. 6B). For example, stemness signatures showed overall negative enrichment in DLBC, BRCA, GBM, and LUSC (Fig. 6B and S6B, Table S10), indicating intrinsic biological heterogeneity between these and other cancer types. Consistently, a negative association between RNAss (a stemness feature extracted from transcriptomic data by machine learning) and survival was also reported in GBM [35]. Also, when we inspected the expression of a number of stemness genes, we only found a few genes, such as SOX2, TWIST1 and CD34, that expressed aberrantly (absolute FC≥2) in more than five cancer types (Fig. S6C), indicating that integrins may function downstream of key stemness factors. Unsupervised clustering grouped proliferation and stemness signatures into one sub-cluster, highlighting an indistinguishable connection between them in tumor evolution (Fig. 6B).

Intriguingly, we found that TME- (except immune cell) and metastasis-associated signatures, including integrin signaling, showed similar enrichment patterns across cancers (Fig. 6B), indicating a role of integrins in TME remodeling that may aid metastasis [5, 6]. EMT empowers cancer cells with morphological plasticity in migrating through barriers, leading to invasion and metastasis [47]. EMT signatures predominately positively enriched in integrinScore high tumors across cancers (Fig. 6B and S6B). It is noteworthy that in bulk RNA-seq data, these EMT signatures should be reflected largely by the stromal compartment within the TME, but not necessarily the epithelial cancer cells. Indeed, we computed the correlation of integrinScore against mRNA expression of many known EMT markers and revealed a preferential positive correlation between integrinScore and mesenchymal markers in the majority of cancer types (Fig. S6D). Especially, this correlation became overwhelmingly significant in most of the adenocarcinomas including STAD, COADREAD, PRAD, and PAAD, indicating intrinsic similarities in metastasis processes in adenocarcinomas originating from different organs. Negative correlations were also found between mesenchymal genes and integrinScore in DLBC, SKCM and KIRC, consistent with the negative enrichment of EMT signatures in the integrinScore-high groups in these cancers (Fig. 6B). The strong correlation between integrinScore and TME/metastasis signatures indicated a predictive value of integrinScore in assessing tumor metastatic potential (Fig. 6B and S6D). To strengthen this idea, we jointly interrogated the TCGA cohort (mainly contained localized primary tumors) and the MET500 dataset (comprised of transcriptomes of 500 metastatic samples from 22 different organs) [48].

The pair-wise comparison revealed that, in most cancer types, integrinScore was significantly higher in metastatic vs. primary tumor samples (Fig. 6C). Notably, primary SKCM samples had higher integrinScore than metastatic samples, which could be explained by the fact that SKCM samples themself were mostly metastatic lymph nodes. Similar integrinScores between primary and metastatic samples in CHOL and THYM was observed, which was consistent with the generally insignificant enrichment of EMT/metastasis signatures in integrinScore high vs. low groups in these cancers (Fig. 6B).

In summary, these results revealed that integrins are expected to interact with diverse biological pathways centric to proliferation, stemness, EMT, metastasis, and immunity, among others, in a context-dependent manner. And as shown in Fig. 6B, we observed 3 subclusters of cancer types with distinct enrichment patterns of signatures in the integrinScore high group, highlighting an intrinsic difference among integrin interactomes in different cancers.

### 7 Integrins remodel TME and predict immunotherapy response

Immune-associated signatures were repeatedly and differentially enriched in tumors stratified by integrinScore across cancers (Fig. 6A and 6B). To search for details, we performed the following analyses. ***First***, we utilized the ESTIMATE algorithm [49] to calculate the immuneScore (proportion of immune cells), stromalScore (proportion of stromal cells), and ESTIMATEScore (proportion of nontumor components) for all TCGA samples. We found that integrinScore was strongly associated with the three scores in most of the cancers (Fig. S7A). Among them, LAML (ρ = 0.70) and TGCT (ρ = -0.76) represented two cancer types with the most positive and negative correlation with integrinScore, respectively (Fig. S7A; Table S11), corroborating the positive and negative enrichment of immune signatures in integrinScore high group in LAML and TGCT, respectively (Fig. 6A and 6B). ***Next***, we utilized the CIBERSORT algorithm to quantify the relative abundance of 22 immune subsets in each tumor sample. As expected, infiltration of different immune cells was all significantly correlated with integrinScore, with heterogeneity in both the correlation direction and strength in a tumor-dependent pattern (Fig. 7A and S7B; Table S11). Importantly, integrinScore generally negatively correlated with antitumor subsets across cancers, including CD8 T cells, activated NK cells, and memory B cells, while positively correlated with immunosuppressive or inactivated subsets, including M2/M0 macrophage and neutrophils (Fig. S7B). To further strengthen our results, we analyzed association between integrinScore and proportion of distinct tumor-infiltrating lymphocytes (TILs) in several PRAD single cell RNA-seq (scRNA-seq) datasets. After annotating the cell identity for each population by commonly-used lineage markers, we calculated the averaged integrinScore for each sample by taking the single cells as a whole. Consistently, we observed negative correlation between integrinScore and the CD8^+^ T cell abundance in two datasets (Fig. S7C, although the p value in GSE141445 dataset was slightly greater than 0.05 due to a small sample size). Expectedly, the correlation between integrinScore and NK proportion in the two datasets was non-significant, in line with the TCGA-PRAD cohort result (Fig. S7C). Antitumor immune response was a series of 7 stepwise events called the cancer-immunity cycle [50]. Based on the relative activity score of each immune steps in the TIP database [50], we computed the correlation between integrinScore and immune steps. Overall, integrinScore significantly correlated with every step in at least four cancers (e.g., step2; Fig. 7B and S7D; Table S11). Among these steps, integrinScore high tumors in 12 cancer types exhibited higher activity in step 1 (release of cancer cell antigens) but lower activity in step 3 (cancer antigen presentation) in the majority of cancers (Fig. S7D), indicating a defect in the antigen presentation process in integrinScore high groups in these cancers. Importantly, integrinScore negatively correlated with the activity of step 7 (killing of cancer cells) in six cancer types, with no positive correlation observed in any other cancers (Fig. S7B and S7D; Table S11). To further dissect the underlying link between integrinScore and tumor immunity, we computed their correlation with the expression of an array of well-known immunomodulators [33]. In line with heterogeneous associations between integrinScore and immune cell components as well as immune cycle steps, we also found that integrinScore showed different correlations with them across cancers (Fig. 7C). Utilizing unsupervised clustering based on correlation, three patterns emerged (negative-, positive- and mix-correlated), regardless of the functional category of immunomodulators (Fig. 7C). Globally, TGCT, HNSC, SKCM, CHOL and SARC were in negative group, consistent with reversal correlations between integrinScore and immune signatures, as well as immune steps, in these cancers (Fig. 6A, 6B, 7B and 7C). Particularly, we observed a negative correlation between some well-known immunotherapeutic targets (i.e., PD1, PD-L1, and CTLA4) and integrinScore in TGCT, HNSC, CESC, LUAD, KIRC, SKCM, CHOL, and SARC. In support, DEG analysis revealed the down-regulation of PD1, PD-L1, and CTLA4 in integrinScore high groups in those cancers (Fig. S7E).

**Figure 7.**
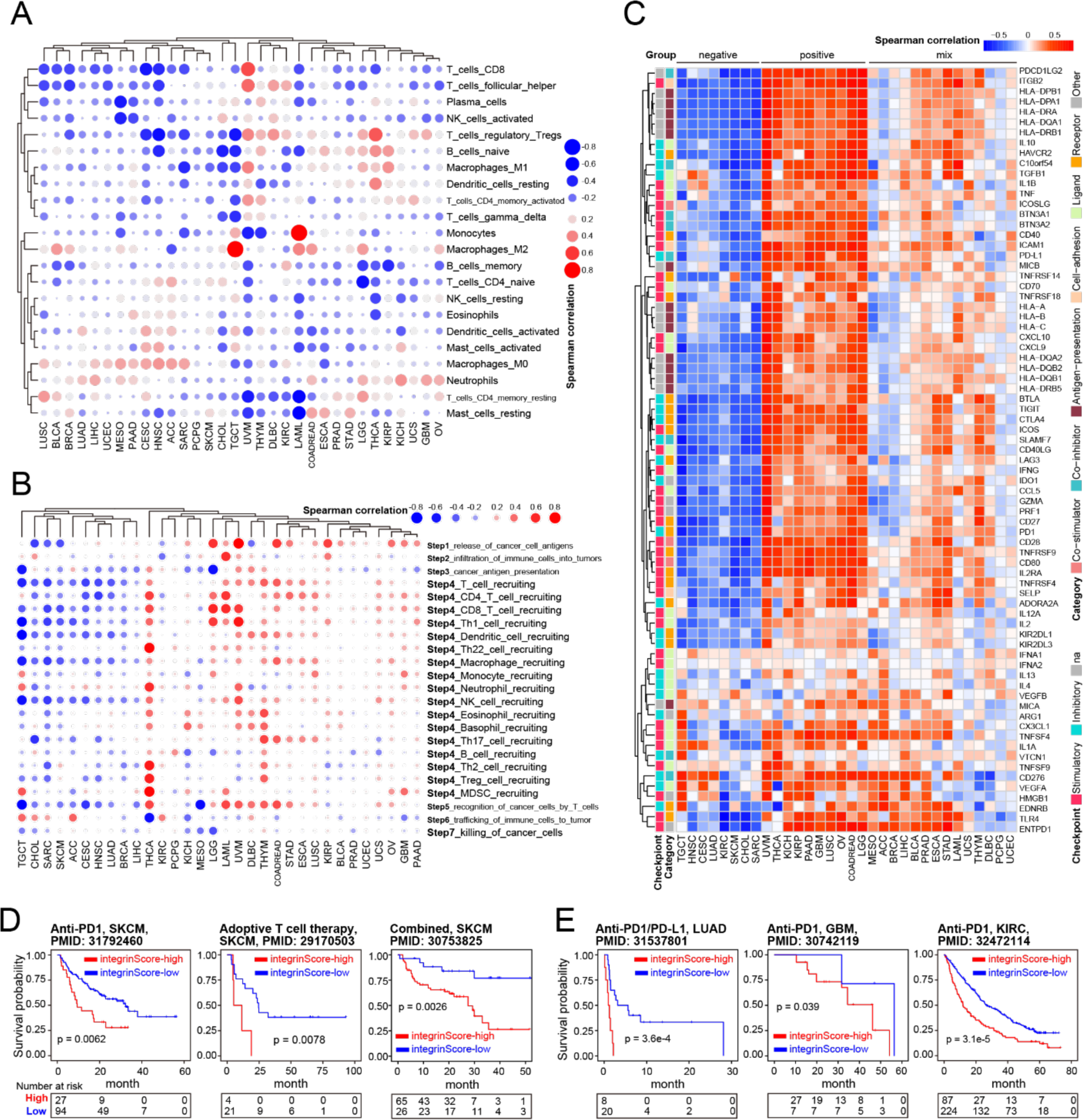
IntegrinScore correlates with immune features in cancer. **A.** Spearman correlation of integrinScore with relative abundance of different types of immune cells across cancers. **B.** Spearman correlation of integrinScore with relative activity of seven immune steps across cancers. Notably, step4 (trafficking of immune cells to tumors) involves various kinds of cells and is showed by cell type. **C.** Heatmap showing Spearman correlation of integrinScore with ICB markers across cancers. Red and bule colors denote positive and negative correlations, respectively. Distinct types of ICB markers are labeled with different colors. Three subgroups are identified by the unsupervised clustering method. **D** and **E**. Kaplan-Meier plots showing integrinScore can predict patient survival after immune therapies in SKCM (**D**) and three other cancers (**E**). For **A** and **B**, blue and red dots denote positive and negative correlations, respectively. The size of dots indicates the absolute correlation coefficient.

The intrinsic link between integrinScore and the immune landscape provided a rationale to ask whether integrinScore can serve as a tool to predict patients’ outcomes to ICB. In three independent SKCM cohorts treated with anti-PD1 [51], adoptive T cell therapy [52], or combined therapy [53], patients were stratified by integrinScore, and we found that patients in the integrinScore low group had significantly longer survival time (Fig. 7D) and a higher percentage of beneficial response to ICB. The percentage of responders in integrinScore high group was 33.3% vs. 42.6%, 0 vs. 47.6% and 44.6% vs. 76.9% compared with that in integrinScore low group in anti-PD1, adoptive T cell, and combined therapy cohort, respectively (Fig. S7F). Similar findings were observed in other three independent cohorts of anti-PD1/PD-L1 in LUAD (n = 28, Fig. 7E and S7G) [54], anti-PD1 in GBM (n = 34, Fig. 7E) [55], and anti-PD1 in KIRC (n = 311, Fig. 7E) [56]. Although differences in response rate between integrinScore high and low groups across some cohorts were not statistically significant due to a limited number of cases, the rate of beneficial response in the integrinScore low group consistently tended to be higher in these cohorts (Fig. S7F and S7G). Collectively, our data established integrinScore, which integrated the interplay between integrins and the immune environment, as a potential biomarker for predicting treatment response to diverse immunotherapies.

### 8 Identification of compounds potentially targeting the integrin family

We next employed the Connectivity Map (CMap) method to search for candidate compounds that might target key signaling pathways intertwined with the integrin family. The CMap is a gene expression profile driven approach designed to uncover associations among genes, chemicals, or other biological perturbations [57]. By a stringent cutoff as having positive or negative correlation with integrinScore in at least 10 cancer types, we identified in total 283 compounds, of which 123 were positively and 160 were negatively correlated (Fig. 8A, Table S12). Mechanism of action (MOA) analysis of all correlated compounds revealed 340 MOAs shared by the above compounds (one compound could have multiple MOAs, Table S12). Topoisomerase inhibitor, CDK inhibitor, and HDAC inhibitor represented the top three MOAs being shared by compounds (Fig. S8A). For the top 20 positively correlated compounds, we observed several of them were TME related, including linifanib (VEGFR inhibitor), SU-11652 (FGFR inhibitor), and PF-562271 (angiogenesis inhibitor). Notably, AKT is downstream of integrin activation [4], and we observed that A-443644 (AKT inhibitor) positively correlated with integrinScore. These results collectively indicated that targeting tumor TME-remodeling and aggressiveness-promoting pathways may represent a strategy to treat cancers driven by overrepresentation of integrin genes. For the top 20 negatively correlated compounds, we found some of them were metabolism-related, including ALW-II-49-7 (glutamate receptor inhibitor), tyrphostin-AG-1295 (dipeptidyl peptidase inhibitor), and sitagliptin (HMGCR inhibitor), implying that these compounds could be used to modulate integrin expression. Targeted gene analysis uncovered 480 unique drug-target genes shared by the abovementioned compounds, with TOP2A, CHRM1/2/3/4, and CDK1/2 being the top ones (Fig. S8B, Table S12).

**Figure 8.**
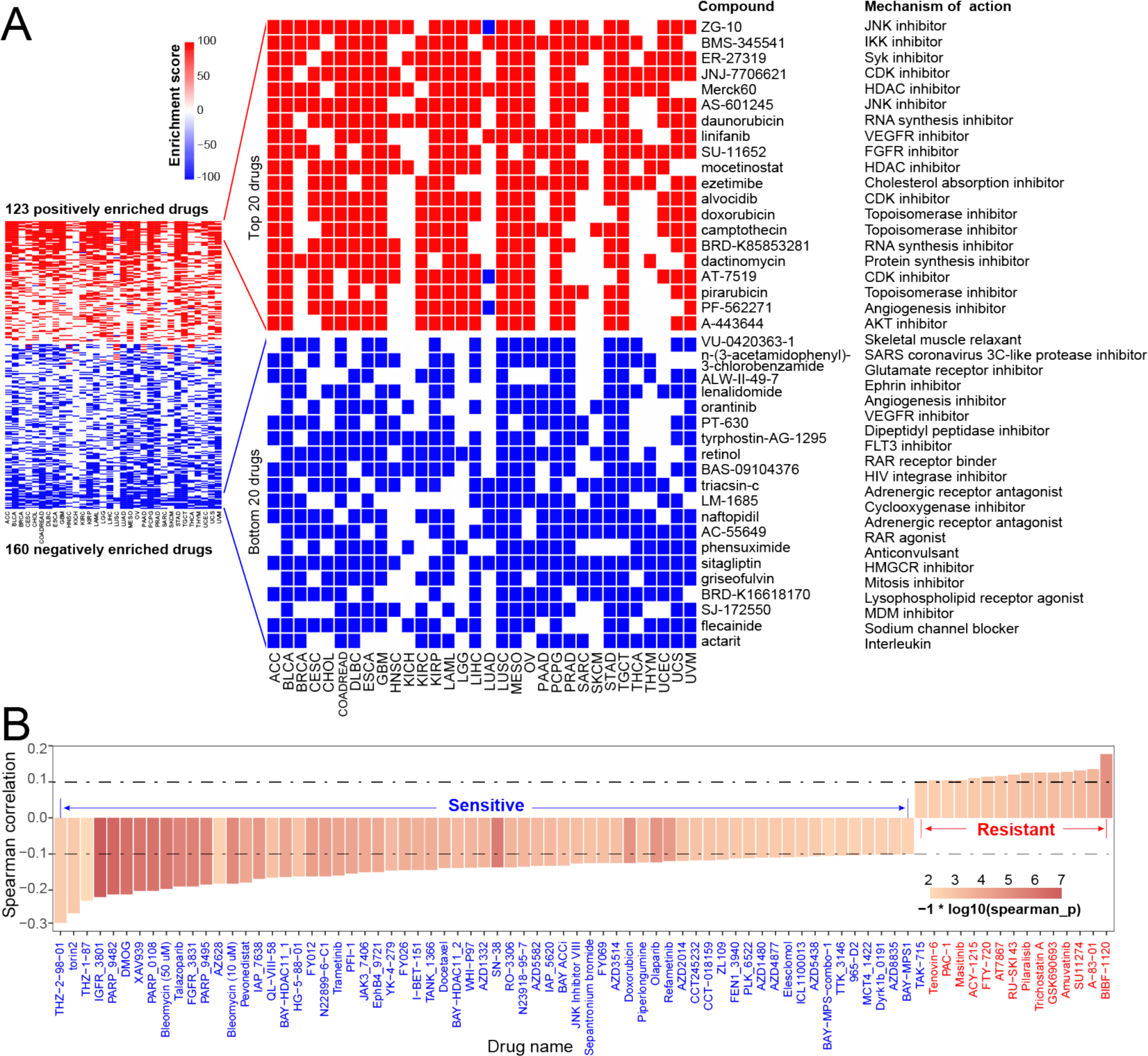
Correlation of integrinScore with drugs. **A.** Heatmap showing the enrichment score of all 283 (left) significantly enriched and extreme 40 compounds (middle) against integrinScore across cancers by the CMap analysis. Red and bule colors denote positive and negative enrichment, respectively. MOA for each compound are listed on the right. **B.** Spearman correlation of integrinScore with IC50 of different compounds reported in the GDSC database. Pairwise results are filtered by absolute correlation coefficient > 0.1 and p < 0.05. Compounds showing positive and negative correlation with integrinScore are considered as resistant (red) and sensitive (blue) drugs.

Alternatively, utilizing the genomics of drug sensitivity in cancer (GDSC) database [58], we calculated the correlation between integrinScore and IC50 for each drug and identified 80 significant correlations (Fig. 8B, Table S13). Specifically, the IC50 values of 15 drugs were positively correlated with integrinScore (indicating drug resistance), whereas 65 drugs were negatively correlated with integrinScore (indicating drug sensitivity), suggesting potential therapeutic strategies to treat integrinScore high tumors with these sensitive drugs. Targeted pathway and gene analysis revealed that genome integrity, kinases, RTK signaling, and PARP1/2 and IAP represented the most affected pathways (Fig. S8C) and genes (Fig. S8D), respectively. PI3K, MAPK, and ERK are downstream of integrin activation [4], we observed that Torin2 and AZD8835 (both targeting PI3K), AZ628 and Refametinib (both targeting ERK/MAPK) negatively correlated with integrinScore, implying an antagonistic role of these compounds against integrin signaling. Interestingly, 33.3% of targeted genes identified in GDSC overlapped with CMap analysis (Fig. S8E), providing an array of actionable targets for interfering aberrant integrin activities.

To further biologically testing these compounds, we chose Torin2 (a potent and selective ATP- competitive mTOR inhibitor) and XAV939 (a Wnt/β-Catenin inhibitor) for experimental validation. Using AR+ LNCaP cell line as a prostate cancer (PCa) model, we showed that Torin (Fig. S8F) and XAV939 (Fig. S8G) treatment for 3 days disturbed the expression of 10/12 and 6/12 integrin genes, respectively. These results provided strong evidence that our identified drugs can modulate the expression pattern of integrin family. Consequently, treatment by these two drugs resulted in significant inhibition of PCa cell proliferation, with Torin2 being more toxic (Fig. S8H). such as AKT and MAKP including p38 and ERK are often downstream of [PMID: 34715893]. Next, we checked the activities of mitotic pathways (i.e., PI3K/AKT, ERK, p38-MAPK) downstream of integrin signaling by Western bolt. Results indicated that not only the total protein level of, but also the activated phosphorylated form, of these pathways were markedly suppressed by Torin2 treatment (Fig. S8I). Collectively, we identified a repertoire of compounds potentially altering integrin gene expression and thus may constitute alternative strategies to targeting cancers harboring aberrant integrin activities.

### 9 Construction of a user-friendly integrin database

To publicly share our findings, we developed an open resource tool called Pan-cancer Integrin Explorer (http://computationalbiology.cn/PIExplorer). Our tool included nine function modules that allow world-wide users to browse and download the majority of integrin-related data presented in this study (Fig. 9A). For most of modules, results were displayed as tables after users submitted their requests, except “Survival”, “Genome” and “Expression” modules in which figures were shown and available for download directly (Fig. 9B and 9C). All tables collected in distinct modules were accessible in the “Download” function (Fig. 9D). Particularly, we also built an “Exploration” module, enabling users to calculate integrinScore for their own samples based on their transcriptome data (Fig. 9E). Therefore, our PIExplorer is a valuable resource for biomedical community to systematically conduct integrin gene analysis across human cancer types.

**Figure 9.**
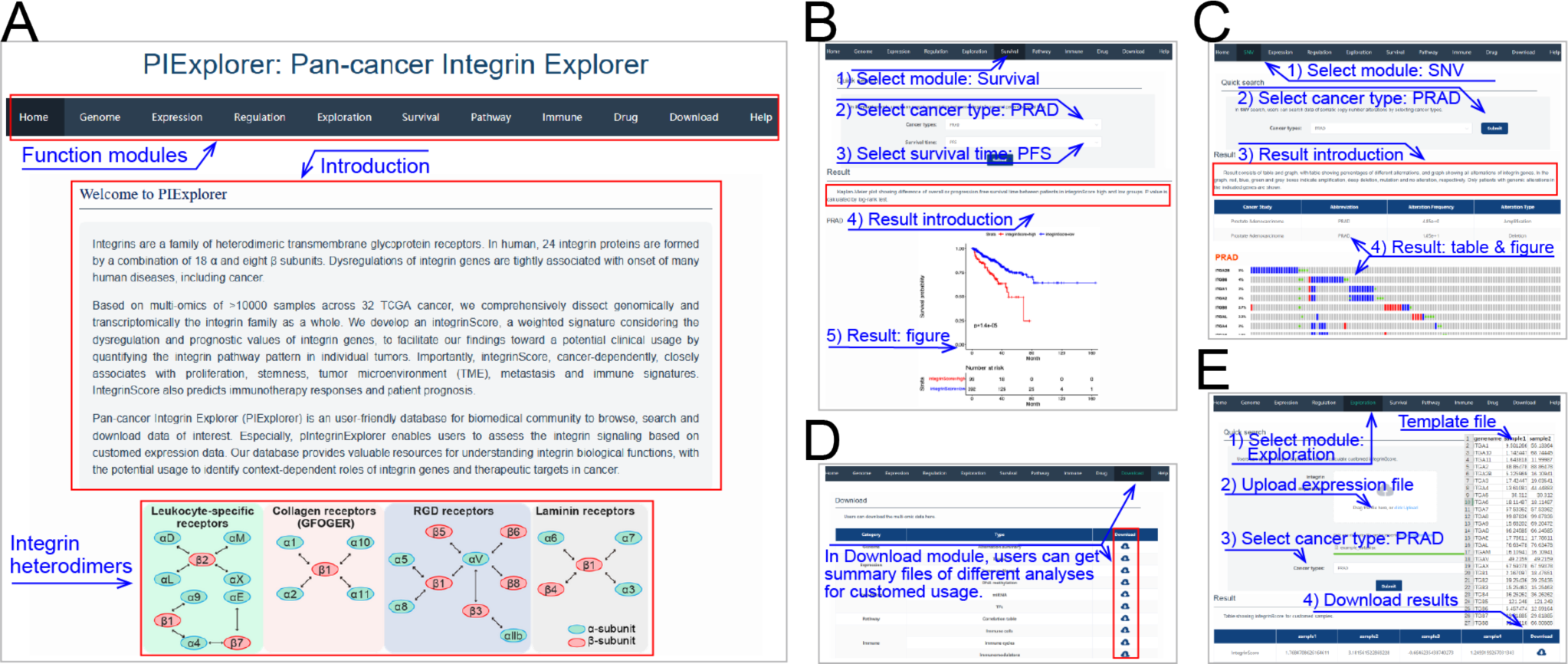
Screenshots for PIExplore database usage. **A.** Screenshots for the homepage of PIExplore website. **B-E**. Usage instructions for “Survival”, “SNV”, “Download”, and “Exploration” modules. Red boxes and blue lines indicate highlighted contents and explanations, respectively.

## DISCUSSION

Integrins are heterodimeric transmembrane receptors that regulate cellular growth, proliferation, and migration via transducing bidirectional biochemical signals between cells [1, 4, 8]. Evidence has implicated integrin in cancer biology mainly through studies of individual genes, yet a comprehensive characterization of the integrin family at multiple genetic levels and associated signatures at the pan-cancer level is lacking. Here, we interrogate TCGA data, including genomic alteration, transcriptomic abnormality (as well as the underlying molecular mechanism such as genetic and epigenetic regulation), and clinical outcome, to systemically profile the integrin pathway in nearly 10,000 tumors of 32 cancer types. Coupled with validation in multiple external cohorts, we unravel not only extensive and context-dependent dysregulation patterns, but also cancer-specific interactomes, of integrins across distinct cancers, providing a global picture of the functional heterogeneity of integrins in human cancers.

We found that although integrins display a very low alteration frequency at individual gene level, they constitute collectively a frequently altered pathway at the population level, indicating involvement in tumorigenesis. In support, studies have shown that T188I mutation in ITGB1 contributes to neoplasia by over-activating ERK/MAPK signaling [59], and mutations in ITGA2 are associated with an increased risk of prostate, oral, and breast cancer [60]. Interestingly, we observe a sexual dimorphism in integrins alteration among female- and male-specific cancers, in that amplification frequency in female cancers is significantly higher than that in male cancers (Fig. 1C). This would suggest a potential relationship between integrins and sex-hormones. Consistently, there are reports that estrogen or androgen can directly bind the regulatory elements of integrin genes to modulate their expression, causing changes in cancer cell behaviors [61–64]. Our findings provide a rationale for considering gender difference in applying integrin-targeted therapies in these particular cancer types.

Compared to the relatively limited number of genomic alterations, transcriptional dysregulation of integrins genes is much more predominant. Changes in the expression of certain integrins have been documented to be linked to tumorigenesis in multiple organs [4, 6, 65]. Here, we show that the majority of integrin genes displaying misexpression are associated with patient outcomes in a context-dependent manner (Fig. 4D and S4). For example, in terms of prognosis analysis, certain integrins may play a protective role in one cancer but an oncogenic role in another cancer. *Vice versa*, in given cancer types, the majority of dysregulated integrins may function as tumor suppressors or promoters. Therefore, our findings highlight an intricate landscape of functional heterogeneity of integrin genes among distinct cancers, which should be considered when we develop and apply integrin-targeted therapies.

Misexpression, but not the genomic alterations, of integrin genes, successfully classify the patient cohorts into 4 clusters exhibiting distinct molecular and clinical features (Fig. 2 and 4). By analyzing genetic (i.e., CNV and TFs) and epigenetic (i.e., DNA methylation and miRNA) mechanisms, we show that dysregulation of integrin genes in cancer is regulated at versatile levels that, in sum, are emerged as a sophisticated regulatory network, expanding our understanding of this pathway compared to previous individual gene-centered studies. Taking the prognostic value and the expression pattern of dysregulated integrin genes in a given cancer type into account, we construct integrinScore to measure an overall integrin signaling pattern in individual tumor samples, with an aim to translate such integrin dysregulation-driven biology toward a potential clinical use. The integrinScore separates each cancer cohort into two groups, with the integrinScore high group being more aggressive. Importantly, integrinScore tightly correlates with numerous predefined genomic, transcriptomic, epigenetic, and clinical classifications, demonstrating utility and generality. Molecularly, we observe that integrinScore cancer-dependently associates with cancer hallmarks and other signatures, including proliferation, stemness, TME/metastasis, and immune features (Fig. 6 and 7). Generally, tumors originate from the same tissue trend to have similar gene expression and regulatory networks [37, 66]. To our surprise, we find the patterns of integrin expression and interactome are obviously different among KIRC, KIRP, and KICH (Fig. 2A, 6A, and 6B). Although few reports have revealed detailed molecule differences among these three kidney cancers, they do differ in biology [67]. Our data imply that, compared to other well-studied oncogenic pathways, integrin signaling may represent a better biomarker to separate these three kidney cancers.

Metastasis is a leading cause of patient death. Here, we surprisingly find that integrinScores are overwhelmingly higher in metastatic vs. primary tumors, consistent with an intrinsic role of integrins in modulating cell motility via downstream functional signaling such as focal adhesion (FAK), Src family kinase (SFK) and phosphoinositide 3-OH kinase (PI3K). Metastatic tumors are usually treatment-resistant. Based on the close relationship between TME and integrinScore, we demonstrate that integrinScore can be used to predict a patient’s response and outcome to ICB. This result could also indicate that a combination of ICB with an integrin-targeted regimen might be beneficial, and there are ongoing clinical trials testing this concept [1, 4, 5]. With an aim to find possible tools that interfere with integrin signaling, we perform pan-cancer analyses based on CMap and GDSC and identify a list of compounds, of which many are targeting downstream effectors of integrins (e.g., AKT1/2/3, FGFR1/2/3, MTOR, and others). We also provide experimental evidence that some of the top hits indeed impact the expression pattern of integrin family, and thus proliferation, in cancer cells, offering a new avenue of combining these potential drugs with ICB to achieve better therapeutic outcomes {Xiong, 2021 #4}.

Collectively, our findings highlight the integrin family as the potential vulnerability for treating aggressive tumors. To facilitate the spreading of our findings, we build an interactive website PIExplorer, to allow the biomedical community to freely check data of interest and evaluate integrin signaling by their own data. There are limitations to our study. First, our results are mainly based on *in silico* analysis, further large-scale experimental validations in different cancer-specific contents are needed. It is noteworthy that we do confirm our results in several other non-TCGA datasets, indicative of the validity of our analytical pipeline. Second, although integrinScore efficiently stratify and predict patients’ immunotherapeutic response at the population level, the model will conceivably suffer limitation when applied to individual patients or a small cohort. This limitation can be reconciled by the combinational use of integrinScore together with other biomarkers to render an accurate stratification. Third, integrin proteins function through heterodimers, our current scoring model does not consider the pairing due to an extreme expression heterogeneity among integrin genes, but an algorithm that weights heterodimers combination based on the transcriptome of 26 integrin genes is expected to further improve the statistical ability of our model. Forth, due to the limited studies of immunotherapies across cancers, we only observe an association between integrinScore and response/outcome in several cohorts (mainly SKCM), clinical pre-application of integrinScore in other cancer types requires more investigation.

## METHODS AND MATERIALS

### Datasets and cohorts

Expression data of mRNA and miRNA, DNA methylation data, and clinical data for TCGA samples were downloaded from the UCSC Xena (https://xenabrowser.net/). Mutation data for TCGA samples were collected from cBioPortal [16]. Expression profiles for validation of the correlation between integrin genes and TFs (Fig. S3K and S3L) were downloaded from GEO (GSE157548) and the CGGA database (http://www.cgga.org.cn/). Transcriptomic and clinical data for validation of integrinScore prognostic value (Fig. S5B) were downloaded from the ICGC data portal (https://dcc.icgc.org/). Expression data for metastatic cancer were collected from the MET500 cohort (https://met500.med.umich.edu/). Expression and clinical data for immunotherapies were downloaded from GEO or collected from the Tumor Immunotherapy Gene Expression Resource portal (tiger.canceromics.org). Known subtypes or classifications of human cancers were collected from original publications as described in the Result section. TCGA tumor type abbreviation codes are as follows: ACC, adrenocortical carcinoma; BLCA, bladder urothelial carcinoma; BRCA, breast invasive carcinoma; CESC, cervical squamous cell carcinoma and endocervical adenocarcinoma; CHOL, cholangiocarcinoma; COAD, colon adenocarcinoma; DLBC, diffuse large B-cell lymphoma; ESCA, oesophageal carcinoma; GBM, glioblastoma multiforme; HNSC, head and neck squamous cell carcinoma; KICH, chromophobe renal cell carcinoma; KIRC, clear cell renal clear cell carcinoma; KIRP, papillary renal cell carcinoma; LAML, acute myeloid leukaemia; LGG, lower-grade glioma; LIHC, hepatocellular carcinoma; LUAD, lung adenocarcinoma; LUSC, lung squamous cell carcinoma; MESO, mesothelioma; OV, ovarian serous adenocarcinoma; PAAD, pancreatic adenocarcinoma; PCPG, phaeochromocytoma and paraganglioma; PRAD, prostate adenocarcinoma; READ, rectal adenocarcinoma; SARC, adult soft tissue sarcoma; SKCM, cutaneous melanoma; STAD, stomach adenocarcinoma; TGCT, testicular germ cell tumour; THCA, thyroid carcinoma; THYM, thymoma; UCEC, uterine corpus endometrial carcinoma; UCS, uterine carcinosarcoma; UVM, uveal melanoma.

### Integrin gene expression analysis

We obtained the normalized gene expression data (count and TPM: Transcripts Per Kilobase of exon model per Million mapped reads) for TCGA samples from the UCSC Xena. For DEG analysis, gene counts for samples were first classified into different groups, then the DESeq2 package [68] was utilized to estimate fold change (FC) of each gene, genes with absolute FC ≥ 2 and false discovery rate (FDR) < 0.05 were identified as DEGs. For other analysis, including multiple association analysis, survival analysis and unsupervised clustering analysis, gene TPM values were utilized (Fig. 2C, 3A, 3D, 3H, 3K, 4D, S2A-C, S2E-H, S3A-D, S3G-I, S3L-M, 4B, 4D, 4E, 4G, S4).

### DNA methylation analysis

The Illumina Human Methylation 450K for 9639 tumor samples was used for analysis. We mapped all probes to individual genes according to their location on the human genome. Gene promoter was defined as the genomic interval from 1000 bp upstream to 200 bp downstream of the transcription start site. Only probes overlapped with integrin genes promoter were retained for further analysis. For multiple probes overlapping with the same integrin gene, the probe with the highest average beta value was used for the final analysis. Significant correlations were filtered by absolute Spearman’s correlation (ρ) ≥ 0.15 and p < 0.05.

### miRNA and TF analysis

For miRNA analysis, we first collected miRNA with available expression data for TCGA samples and annotated by StarBase [69] and miRTarBase [70] simultaneously. To remove miRNAs with extremely low expression, only miRNAs with normalized expression >1 in more than 500 samples were retained. The correlation between miRNA and integrin genes was calculated by the Spearman method and pairs with absolute ρ ≥ 0.3 and p < 0.05 in more than five cancer types were regarded as significantly correlated. To enhance data quality, 40 significant miRNA- integrin pairs with validation in StarBase or miRTarBase were used for final analysis.

For TF analysis, we first downloaded all TF-target pairs, including 495 TFs and 38183 targets, from the hTFtarget database (http://bioinfo.life.hust.edu.cn/hTFtarget/#!/). The correlation between TFs and target genes was calculated by Spearman and pairs with absolute ρ ≥ 0.2 and p < 0.01 were retained for further analysis. The top 10 TFs with the most integrin target genes were shown (Fig. 3J). For integrated analysis of miRNA and TF (Fig. S3N), we first extracted 27 miRNAs and 11 integrins in the 40 miRNA-integrin pairs. Then, 21 TFs targeting all 11 integrins were identified from the hTFtarget database. Next, associations between the 27 miRNAs and 21 TFs were calculated by miRTarBase. Finally, a reciprocally-regulated network between miRNAs, TFs, and integrins was constructed by Cytoscape software.

### Survival analysis

The survival differences between different groups (e.g. patients with/without integrin gene alteration, Fig. S1B-S1E; in different clusters, Fig. 2B) were estimated by Kaplan-Meier curve, and p < 0.05 was considered as statistically significant. For continuous variables like gene expression or integrinScore, the ‘survcutpoint’ function of “survminer” package was first applied to determine the optimal cutoff for maximum rank statistic, then the “survival” R package was utilized to compare survival differences between high and low groups.

### IntegrinScore computation

We constructed integrinScore for each sample in a way used in our recent study [38]. Briefly, we first performed survival analysis for integrin genes across 32 cancer types. We utilized both survival outcomes (OS and PFS) in a cancer-dependent manner in survival analysis, as recommended previously {Liu, 2018 #85} . The oncogenic integrin gene was identified as when patients had higher gene expression, their survival time was significantly shorter; while the protective integrin gene was identified by the opposite trend. Then, 26 integrin gene expressions for each sample were extracted and normalized. And the integrinScore was calculated based on a linear combination of normalized expression values of the 26 integrin genes, of which oncogenic and protective integrin genes contributed positively and negatively to integrinScore, respectively:

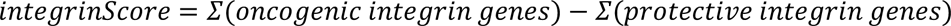

### Biological pathway/signature analysis

We identified biological pathways/signatures associated with integrinScore by the pre-ranked module embedded in gene set enrichment analysis (GSEA) [71]. After stratifying patients into integrinScore high and low groups, DEGs were identified and all genes were ranked by FC. Then, cancer hallmarks and curated signatures (mainly from c2-all which contained 7233 gene- sets) were downloaded from the GSEA website were enriched against ranked gene lists by clusterProfiler package [72], and signatures with FDR < 0.1 were considered statistically significant. For the proliferation related signature in Fig. 4B, the relative activity score of four signatures was estimated by gene set variation analysis (GSVA) package [36] in the R environment.

### Immune cell, cycle and modulator analysis

The estimation of stromal and immune cells in malignant tumor tissues using the expression data (ESTIMATE) algorithm was used to score for tumor purity, the level of stromal cells present, and immune cell infiltration in tumor tissues based on expression data [49]. The CIBERSORT is a deconvolution tool to estimate the abundances of immune cell types in a mixed cell population using gene expression data [73]. We downloaded CIBERSORT results for TCGA pan-cancer samples from the GDC PanCan portal (https://gdc.cancer.gov/about-data/publications/panimmune). A previous study has conceptualized the anticancer immune response as seven sequential steps: (i) release of cancer cell antigens; (ii) cancer antigen presentation; (iii) priming and activation; (iv) trafficking of immune cells to tumors; (v) infiltration of immune cells into tumors; (vi) recognition of cancer cells by T cells; and (vii) killing of cancer cells. In aggregate, these seven steps were referred to as the cancer immunity cycle [50]. We downloaded the relative activity score of seven steps for TCGA pan-cancer samples from the TIP portal (http://biocc.hrbmu.edu.cn/TIP/). An array of immunomodulators were obtained from TCGA immune response working group [33]. Spearman’s correlation coefficients between integrinScore and immune cells/cycles/ modulators were calculated for distinct cancers.

### Drug analysis

For CMap analysis, we first performed DEG analysis for samples stratified by integrinScore of each cancer type. Then top 150 upregulated genes (integrinScore high vs. low group) were subjected to the query module of the CMap online tool (https://clue.io/). Compounds with enrichment scores > 95 or < -95 in at least ten cancer types were considered as significantly positive or negative compounds, respectively. For GDSC analysis, we first calculated integrinScore for each cell line according to distinct cancer types. Then, the Spearman correlation between integrinScore and IC50 of different drugs was calculated and pairs with absolute ρ ≥ 0.1 and p < 0.05 were considered significantly correlated.

### Drug treatment, MTT assay, Quantitative RT-PCR, and Western blot analysis

For all drug treatment experiments, drugs were dissolved in dimethyl sulfoxide (DMSO) separately. The effects of single drugs were pre-determined and doses lower than IC50 were used. Torin, 5’Aza (XXX), Enzalutamide (XXX), and XAV939 (XXX) were purchased from TOPScience. For MTT assays, cells were trypsinized, counted and seeded in 96-well plates, then, after drug treatment for 3-4 days, MTT was added at a concentration of 0.5 mg/ml for 3 h at 37 °C. The medium was then removed and 0.2 ml/well of acidic isopropyl alcohol (0.04M HCl in absolute isopropyl alcohol) was added. The absorbance of the converted dye was measured at 570 nm using a Synergy II spectrophotometer (Biotek). Quantitative RT-PCR was performed using the iQ SYBR Green supermix (BioRad) on a 7900HT Fast Real-Time PCR System (Applied Biosystems). The primers used in this study were listed in Table S14. Normally, the housekeeping gene GAPDH or β-actin was used as internal control for gene expression normalization. For western blot analysis, antibodies against p-AKT, AKT1/2/3, p-ERK1/2, ERK1/2, p-p38, p38, and β-Tubulin were used (Table S14).

### Database construction

We constructed the PIExplorer database by the standard MVC (Model-View-Controller) pattern, which consists of a server side and a client side. The front-end was built by Vue and Element UI. The backend was encoded by Java, and raw data was stored and managed by the MySQL, with Spring Boot, Mybatisplus and Redis providing interface services. The outputs of database are tables and/or figures.

### Statistical analysis

All statistical analyses in our study were performed by R 4.1.0. Specifically, analysis of methylation, miRNA, and TFs, as well as the association between integrinScore and selected signatures, cancer immunity cycle, 22 immune cell subpopulations, and marker genes, were performed by Spearman analysis (Fig. 3D, 3H, 3K, S5C, S6D, 7A-7C, and S7A). Quantitative data fitting normal distribution were compared by t-test; otherwise, the Wilcoxon (for two groups) or Kruskal–Wallis (for more than two groups) test was used. Chi-square or Fisher’s exact test was performed to compare differences between categorical variables. Heatmaps showing Spearman ρ, FC, and p value (Fig. 2A, S2H, 3D, 3H, 3K, 4D, S4, 6A, 6B, S6C, S6D, 7A, 7B, and S7A) were visualized by TBTools [74].

## AUTHOR CONTRIBUTIONS

**Cheng Zou**: Conceptualization, Methodology, Software, Validation, Formal analysis, Investigation, Data Curation, Writing - Original Draft, Visualization, Funding acquisition.

**Jinwei Zhu**: Methodology, Software, Formal analysis, Investigation, Data Curation.

**Jiangling Xiong**: Investigation, Data Curation, Experimental Validation.

**Yu Tian**: Experimental Validation.

**Yousong Peng**: Methodology, Software, Resources.

**Edwin Cheung**: Methodology, Software, Resources.

**Dingxiao Zhang**: Conceptualization, Methodology, Writing - Original Draft, Writing - Review & Editing, Visualization, Supervision, Project administration, Funding acquisition.

## COMPETING INTERESTS

The authors declare no competing interests.

## ACKNOWLEDGMENTS

We thank Dr. Ping Fu (College of Biology, Human University) for her technical advice on integrin database construction.

## FUNDING

This work was supported by Grants from the National Natural Science Foundation of China (81972418, DZ), the Hunan Provincial Science Fund for Distinguished Young Scholars (2021JJ10028, DZ), the Shenzhen Natural Science Foundation (JCYJ20220530160410024, DZ), and the Fundamental Research Funds for the Central Universities (DZ). C.Z. was supported, in part, by the Hunan Province Natural Science Foundation (2022JJ40111, CZ) and the Changsha Natural Science Foundation (kq2202182, CZ).

## Notes

### Competing Interest Statement

The authors have declared no competing interest.

## References

[1] Slack RJ, Macdonald SJF, Roper JA, Jenkins RG, Hatley RJD. Emerging therapeutic opportunities for integrin inhibitors. Nat Rev Drug Discov 2022;21:60–78.

[2] Humphries JD, Byron A, Humphries MJ. Integrin ligands at a glance. J Cell Sci 2006;119:3901–3.

[3] Winograd-Katz SE, Fassler R, Geiger B, Legate KR. The integrin adhesome: from genes and proteins to human disease. Nat Rev Mol Cell Biol 2014;15:273–88.

[4] Xiong J, Yan L, Zou C, Wang K, Chen M, Xu B, et al. Integrins regulate stemness in solid tumor: an emerging therapeutic target. J Hematol Oncol 2021;14:177.

[5] Cooper J, Giancotti FG. Integrin Signaling in Cancer: Mechanotransduction, Stemness, Epithelial Plasticity, and Therapeutic Resistance. Cancer Cell 2019;35:347–67.

[6] Hamidi H, Ivaska J. Every step of the way: integrins in cancer progression and metastasis. Nature Reviews Cancer 2018;18:532–47.

[7] Ren B, Yu YP, Tseng GC, Wu C, Chen K, Rao UN, et al. Analysis of integrin alpha7 mutations in prostate cancer, liver cancer, glioblastoma multiforme, and leiomyosarcoma. J Natl Cancer Inst 2007;99:868–80.

[8] Desgrosellier JS, Cheresh DA. Integrins in cancer: biological implications and therapeutic opportunities. Nat Rev Cancer 2010;10:9–22.

[9] Zhang D, Tang DG, Rycaj K. Cancer stem cells: Regulation programs, immunological properties and immunotherapy. Semin Cancer Biol 2018.

[10] Zhang Y, Wang H. Integrin signalling and function in immune cells. Immunology 2012;135:268–75.

[11] Wang H, Lim D, Rudd CE. Immunopathologies linked to integrin signalling. Semin Immunopathol 2010;32:173–82.

[12] Pribila JT, Quale AC, Mueller KL, Shimizu Y. Integrins and T cell-mediated immunity. Annu Rev Immunol 2004;22:157–80.

[13] Busenhart P, Montalban-Arques A, Katkeviciute E, Morsy Y, Van Passen C, Hering L, et al. Inhibition of integrin alphavbeta6 sparks T-cell antitumor response and enhances immune checkpoint blockade therapy in colorectal cancer. J Immunother Cancer 2022;10.

[14] Bagati A, Kumar S, Jiang P, Pyrdol J, Zou AE, Godicelj A, et al. Integrin alphavbeta6-TGFbeta- SOX4 Pathway Drives Immune Evasion in Triple-Negative Breast Cancer. Cancer Cell 2021;39:54–67 e9.

[15] Tomczak K, Czerwinska P, Wiznerowicz M. The Cancer Genome Atlas (TCGA): an immeasurable source of knowledge. Contemp Oncol (Pozn) 2015;19:A68–77.

[16] Gao J, Aksoy BA, Dogrusoz U, Dresdner G, Gross B, Sumer SO, et al. Integrative analysis of complex cancer genomics and clinical profiles using the cBioPortal. Sci Signal 2013;6:pl1.

[17] Lawrence MS, Stojanov P, Polak P, Kryukov GV, Cibulskis K, Sivachenko A, et al. Mutational heterogeneity in cancer and the search for new cancer-associated genes. Nature 2013;499:214–8.

[18] Stange DE, Radlwimmer B, Schubert F, Traub F, Pich A, Toedt G, et al. High-resolution genomic profiling reveals association of chromosomal aberrations on 1q and 16p with histologic and genetic subgroups of invasive breast cancer. Clin Cancer Res 2006;12:345–52.

[19] Group PTC, Calabrese C, Davidson NR, Demircioglu D, Fonseca NA, He Y, et al. Genomic basis for RNA alterations in cancer. Nature 2020;578:129–36.

[20] Beroukhim R, Mermel CH, Porter D, Wei G, Raychaudhuri S, Donovan J, et al. The landscape of somatic copy-number alteration across human cancers. Nature 2010;463:899–905.

[21] Moore LD, Le T, Fan G. DNA methylation and its basic function. Neuropsychopharmacology 2013;38:23–38.

[22] Lee YS, Dutta A. MicroRNAs in cancer. Annu Rev Pathol 2009;4:199–227.

[23] Esquela-Kerscher A, Slack FJ. Oncomirs - microRNAs with a role in cancer. Nat Rev Cancer 2006;6:259–69.

[24] Muller DW, Bosserhoff AK. Integrin beta 3 expression is regulated by let-7a miRNA in malignant melanoma. Oncogene 2008;27:6698–706.

[25] Bronisz A, Rooj AK, Krawczynski K, Peruzzi P, Salinska E, Nakano I, et al. The nuclear DICER- circular RNA complex drives the deregulation of the glioblastoma cell microRNAome. Sci Adv 2020;6.

[26] Bushweller JH. Targeting transcription factors in cancer - from undruggable to reality. Nat Rev Cancer 2019;19:611–24.

[27] Stine ZE, Walton ZE, Altman BJ, Hsieh AL, Dang CV. MYC, Metabolism, and Cancer. Cancer Discov 2015;5:1024–39.

[28] Zhang Q, Liu W, Zhang HM, Xie GY, Miao YR, Xia M, et al. hTFtarget: A Comprehensive Database for Regulations of Human Transcription Factors and Their Targets. Genomics Proteomics Bioinformatics 2020;18:120–8.

[29] Dang CV. MYC on the path to cancer. Cell 2012;149:22–35.

[30] He Y, Xu W, Xiao YT, Huang H, Gu D, Ren S. Targeting signaling pathways in prostate cancer: mechanisms and clinical trials. Signal Transduct Target Ther 2022;7:198.

[31] Okita K, Nakagawa M, Hyenjong H, Ichisaka T, Yamanaka S. Generation of mouse induced pluripotent stem cells without viral vectors. Science 2008;322:949–53.

[32] Zhao Z, Zhang KN, Wang Q, Li G, Zeng F, Zhang Y, et al. Chinese Glioma Genome Atlas (CGGA): A Comprehensive Resource with Functional Genomic Data from Chinese Glioma Patients. Genomics Proteomics Bioinformatics 2021;19:1–12.

[33] Thorsson V, Gibbs DL, Brown SD, Wolf D, Bortone DS, Ou Yang TH, et al. The Immune Landscape of Cancer. Immunity 2018;48:812–30 e14.

[34] Hanahan D. Hallmarks of Cancer: New Dimensions. Cancer Discov 2022;12:31–46.

[35] Malta TM, Sokolov A, Gentles AJ, Burzykowski T, Poisson L, Weinstein JN, et al. Machine Learning Identifies Stemness Features Associated with Oncogenic Dedifferentiation. Cell 2018;173:338–54 e15.

[36] Hanzelmann S, Castelo R, Guinney J. GSVA: gene set variation analysis for microarray and RNA-seq data. BMC Bioinformatics 2013;14:7.

[37] Luo Z, Liu W, Sun P, Wang F, Feng X. Pan-cancer analyses reveal regulation and clinical outcome association of the shelterin complex in cancer. Brief Bioinform 2021;22.

[38] Zou C, He Q, Feng Y, Chen M, Zhang D. A m(6)Avalue predictive of prostate cancer stemness, tumor immune landscape and immunotherapy response. NAR Cancer 2022;4:zcac010.

[39] Bagaev A, Kotlov N, Nomie K, Svekolkin V, Gafurov A, Isaeva O, et al. Conserved pan-cancer microenvironment subtypes predict response to immunotherapy. Cancer Cell 2021;39:845–65 e7.

[40] Zheng S, Cherniack AD, Dewal N, Moffitt RA, Danilova L, Murray BA, et al. Comprehensive Pan-Genomic Characterization of Adrenocortical Carcinoma. Cancer Cell 2016;29:723–36.

[41] Ceccarelli M, Barthel FP, Malta TM, Sabedot TS, Salama SR, Murray BA, et al. Molecular Profiling Reveals Biologically Discrete Subsets and Pathways of Progression in Diffuse Glioma. Cell 2016;164:550–63.

[42] Campbell JD, Yau C, Bowlby R, Liu Y, Brennan K, Fan H, et al. Genomic, Pathway Network, and Immunologic Features Distinguishing Squamous Carcinomas. Cell Rep 2018;23:194–212 e6.

[43] Gabrilovich DI, Ostrand-Rosenberg S, Bronte V. Coordinated regulation of myeloid cells by tumours. Nat Rev Immunol 2012;12:253–68.

[44] Ouzounova M, Lee E, Piranlioglu R, El Andaloussi A, Kolhe R, Demirci MF, et al. Monocytic and granulocytic myeloid derived suppressor cells differentially regulate spatiotemporal tumour plasticity during metastatic cascade. Nat Commun 2017;8:14979.

[45] Moffitt RA, Marayati R, Flate EL, Volmar KE, Loeza SG, Hoadley KA, et al. Virtual microdissection identifies distinct tumor- and stroma-specific subtypes of pancreatic ductal adenocarcinoma. Nat Genet 2015;47:1168–78.

[46] Bailey P, Chang DK, Nones K, Johns AL, Patch AM, Gingras MC, et al. Genomic analyses identify molecular subtypes of pancreatic cancer. Nature 2016;531:47–52.

[47] Hanahan D, Weinberg RA. Hallmarks of cancer: the next generation. Cell 2011;144:646–74.

[48] Robinson DR, Wu YM, Lonigro RJ, Vats P, Cobain E, Everett J, et al. Integrative clinical genomics of metastatic cancer. Nature 2017;548:297–303.

[49] Yoshihara K, Shahmoradgoli M, Martinez E, Vegesna R, Kim H, Torres-Garcia W, et al. Inferring tumour purity and stromal and immune cell admixture from expression data. Nat Commun 2013;4:2612.

[50] Xu L, Deng C, Pang B, Zhang X, Liu W, Liao G, et al. TIP: A Web Server for Resolving Tumor Immunophenotype Profiling. Cancer Res 2018;78:6575–80.

[51] Liu D, Schilling B, Liu D, Sucker A, Livingstone E, Jerby-Arnon L, et al. Integrative molecular and clinical modeling of clinical outcomes to PD1 blockade in patients with metastatic melanoma. Nat Med 2019;25:1916–27.

[52] Lauss M, Donia M, Harbst K, Andersen R, Mitra S, Rosengren F, et al. Mutational and putative neoantigen load predict clinical benefit of adoptive T cell therapy in melanoma. Nat Commun 2017;8:1738.

[53] Gide TN, Quek C, Menzies AM, Tasker AT, Shang P, Holst J, et al. Distinct Immune Cell Populations Define Response to Anti-PD-1 Monotherapy and Anti-PD-1/Anti-CTLA-4 Combined Therapy. Cancer Cell 2019;35:238–55 e6.

[54] Jung H, Kim HS, Kim JY, Sun JM, Ahn JS, Ahn MJ, et al. DNA methylation loss promotes immune evasion of tumours with high mutation and copy number load. Nat Commun 2019;10:4278.

[55] Zhao J, Chen AX, Gartrell RD, Silverman AM, Aparicio L, Chu T, et al. Immune and genomic correlates of response to anti-PD-1 immunotherapy in glioblastoma. Nat Med 2019;25:462–9.

[56] Braun DA, Hou Y, Bakouny Z, Ficial M, Sant’ Angelo M, Forman J, et al. Interplay of somatic alterations and immune infiltration modulates response to PD-1 blockade in advanced clear cell renal cell carcinoma. Nat Med 2020;26:909–18.

[57] Lamb J, Crawford ED, Peck D, Modell JW, Blat IC, Wrobel MJ, et al. The Connectivity Map: using gene-expression signatures to connect small molecules, genes, and disease. Science 2006;313:1929–35.

[58] Yang W, Soares J, Greninger P, Edelman EJ, Lightfoot H, Forbes S, et al. Genomics of Drug Sensitivity in Cancer (GDSC): a resource for therapeutic biomarker discovery in cancer cells. Nucleic Acids Res 2013;41:D955–61.

[59] Evans RD, Perkins VC, Henry A, Stephens PE, Robinson MK, Watt FM. A tumor-associated beta 1 integrin mutation that abrogates epithelial differentiation control. J Cell Biol 2003;160:589–96.

[60] Marthick JR, Dickinson JL. Emerging putative biomarkers: the role of alpha 2 and 6 integrins in susceptibility, treatment, and prognosis. Prostate Cancer 2012;2012:298732.

[61] Singhal H, Greene ME, Tarulli G, Zarnke AL, Bourgo RJ, Laine M, et al. Genomic agonism and phenotypic antagonism between estrogen and progesterone receptors in breast cancer. Sci Adv 2016;2:e1501924.

[62] Asangani IA, Dommeti VL, Wang X, Malik R, Cieslik M, Yang R, et al. Therapeutic targeting of BET bromodomain proteins in castration-resistant prostate cancer. Nature 2014;510:278–82.

[63] Geoghegan IP, Hoey DA, McNamara LM. Estrogen deficiency impairs integrin alpha(v)beta(3)- mediated mechanosensation by osteocytes and alters osteoclastogenic paracrine signalling. Sci Rep 2019;9:4654.

[64] Davis PJ, Mousa SA, Cody V, Tang HY, Lin HY. Small molecule hormone or hormone-like ligands of integrin alphaVbeta3: implications for cancer cell behavior. Horm Cancer 2013;4:335–42.

[65] Seguin L, Desgrosellier JS, Weis SM, Cheresh DA. Integrins and cancer: regulators of cancer stemness, metastasis, and drug resistance. Trends Cell Biol 2015;25:234–40.

[66] Wang X, Ma L, Pei X, Wang H, Tang X, Pei JF, et al. Comprehensive assessment of cellular senescence in the tumor microenvironment. Brief Bioinform 2022;23.

[67] Chen F, Zhang Y, Senbabaoglu Y, Ciriello G, Yang L, Reznik E, et al. Multilevel Genomics-Based Taxonomy of Renal Cell Carcinoma. Cell Rep 2016;14:2476–89.

[68] Love MI, Huber W, Anders S. Moderated estimation of fold change and dispersion for RNA-seq data with DESeq2. Genome Biol 2014;15:550.

[69] Li JH, Liu S, Zhou H, Qu LH, Yang JH. starBase v2.0: decoding miRNA-ceRNA, miRNA-ncRNA and protein-RNA interaction networks from large-scale CLIP-Seq data. Nucleic Acids Res 2014;42:D92–7.

[70] Huang HY, Lin YC, Cui S, Huang Y, Tang Y, Xu J, et al. miRTarBase update 2022: an informative resource for experimentally validated miRNA-target interactions. Nucleic Acids Res 2022;50:D222–D30.

[71] Subramanian A, Tamayo P, Mootha VK, Mukherjee S, Ebert BL, Gillette MA, et al. Gene set enrichment analysis: a knowledge-based approach for interpreting genome-wide expression profiles. Proc Natl Acad Sci U S A 2005;102:15545–50.

[72] Wu T, Hu E, Xu S, Chen M, Guo P, Dai Z, et al. clusterProfiler 4.0: A universal enrichment tool for interpreting omics data. Innovation (Camb) 2021;2:100141.

[73] Newman AM, Liu CL, Green MR, Gentles AJ, Feng W, Xu Y, et al. Robust enumeration of cell subsets from tissue expression profiles. Nat Methods 2015;12:453–7.

[74] Chen C, Chen H, Zhang Y, Thomas HR, Frank MH, He Y, et al. TBtools: An Integrative Toolkit Developed for Interactive Analyses of Big Biological Data. Mol Plant 2020;13:1194–202.

