## Supplementary Info for "Comprehensive characterization of the integrin family across 32 cancer types"

**Cheng Zou et al.**

#### Summary of Supplementary Materials

**Supplementary Figures:**

Related to Fig. S1
Fig. S1 Infrequently altered integrin genes are rarely associated with patients' survival.
Related to Fig. S2
Fig. S2 Expression pattern of integrin genes correlates with patients' survival.
Related to Fig. S3
Fig. S3 Mechanisms underpinning integrin gene dysregulation.
Related to Fig. S4
Fig. S4 Individual integrin gene expression frequently correlated with patients' survival.
Related to Fig. S5
Fig. S5 IntegrinScore correlates with known subtypes with higher aggressiveness.
Related to Fig. S6
Fig. S6 IntegrinScore correlates with cancer hallmarks.
Related to Fig. S7
Fig. S7 IntegrinScore correlates with immune landscape.
Related to Fig. S8
Fig. S8 Representative pathways and genes potentially targeted by identified compounds.

**Supplementary Tables:**

Table S1. Abbreviation of 32 cancer types in TCGA
Table S2. Alteration Frequency of 26 integrin genes across 32 cancer types in TCGA
Table S3. Fold change and FDR of 26 integrin genes across 15 cancer types
Table S4. Cluster (k=4) and survival information for TCGA samples
Table S5. P value and spearman correlation between DNA methylation and expression of integrin genes in pan-cancer
Table S6. P value and spearman correlation between miRNA and integrin gene expression in pan-cancer
Table S7. P value and spearman correlation between TFs and integrin gene expression in pan-cancer
Table S8. P value for analysis of integrin genes and survival time in pan-cancer
Table S9. Normalized enrichment score (NES) and FDR of 50 hallmarks analyzed by GSEA across 32 cancer types
Table S10. Normalized enrichment score (NES) and FDR of 40 curated signatures analyzed by GSEA across 32 cancer types
Table S11. P value and spearman correlation between integrinScore and immune features in pan-cancer
Table S12. Enrichment score of each compound from the CMap across 32 cancer types
Table S13. P value and spearman correlation between integrinScore and IC50 of different drugs in GDSC
Table S14. List of primers and antibodies used in this study

### Supplementary Figures

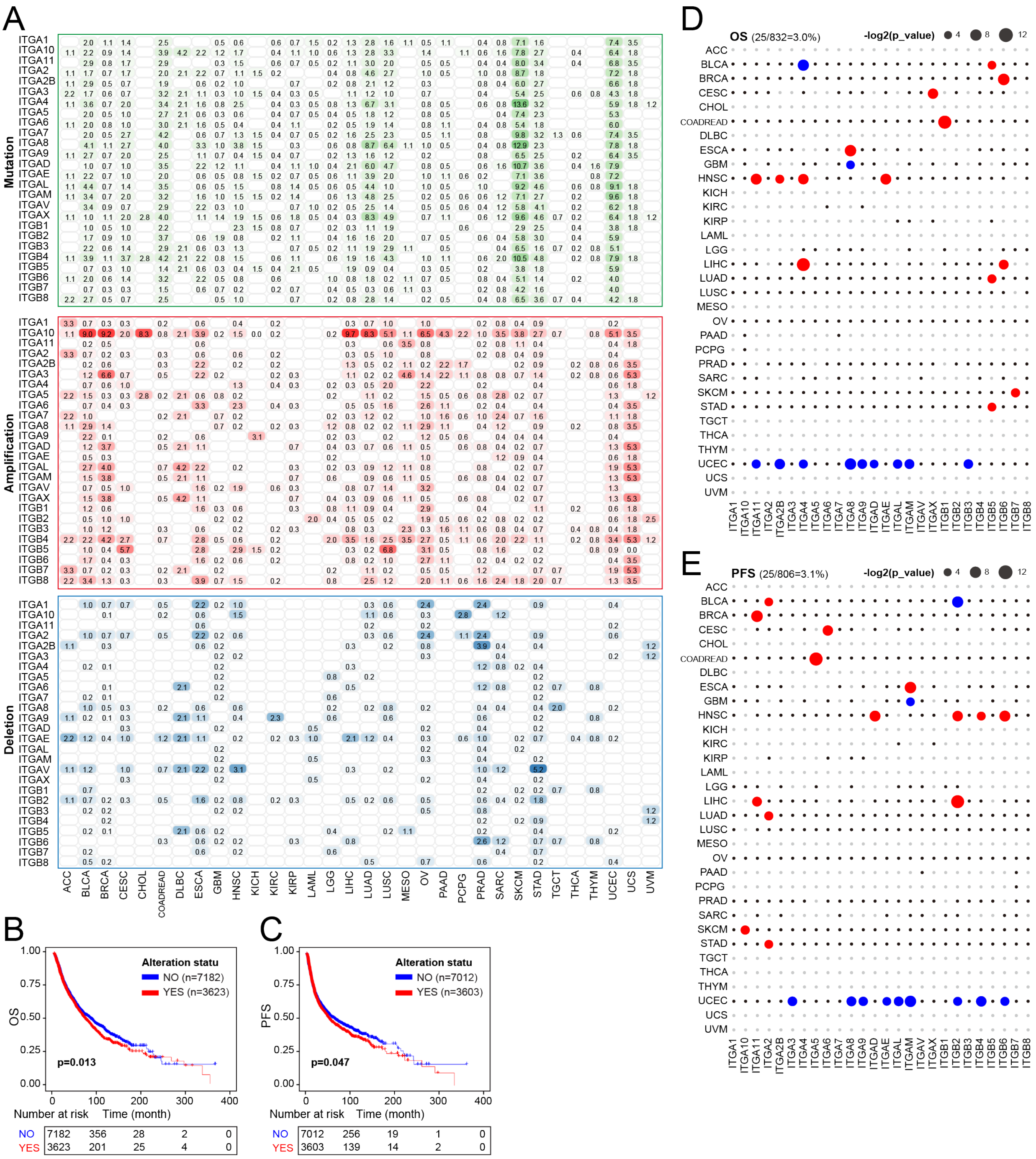


**Fig. S1. Integrin alterations rarely correlate with patient's survival time**

**A.** Distribution of mutation, amplification, and deletion frequencies of 26 integrin genes across cancers. Data shown in the pane are frequency values.

**B and C.** Kaplan-Meier plots show that integrin alterations are associated with shorter overall survival (OS, **B**) and progression free survival (PFS, **C**) time.

**D and E.** Summary of survival analysis results (OS in **D**; PFS in **E**) of integrin genes across cancers. Red, blue and black dots indicate worse, better and unsignificant results, respectively. Alteration groups with <5 samples are not performed survival analysis and denoted as NA (grey dots). The size of dots indicates $-\log_2$(p_values). Data in the parentheses (top left) denote the rate of alterations associated with survival.

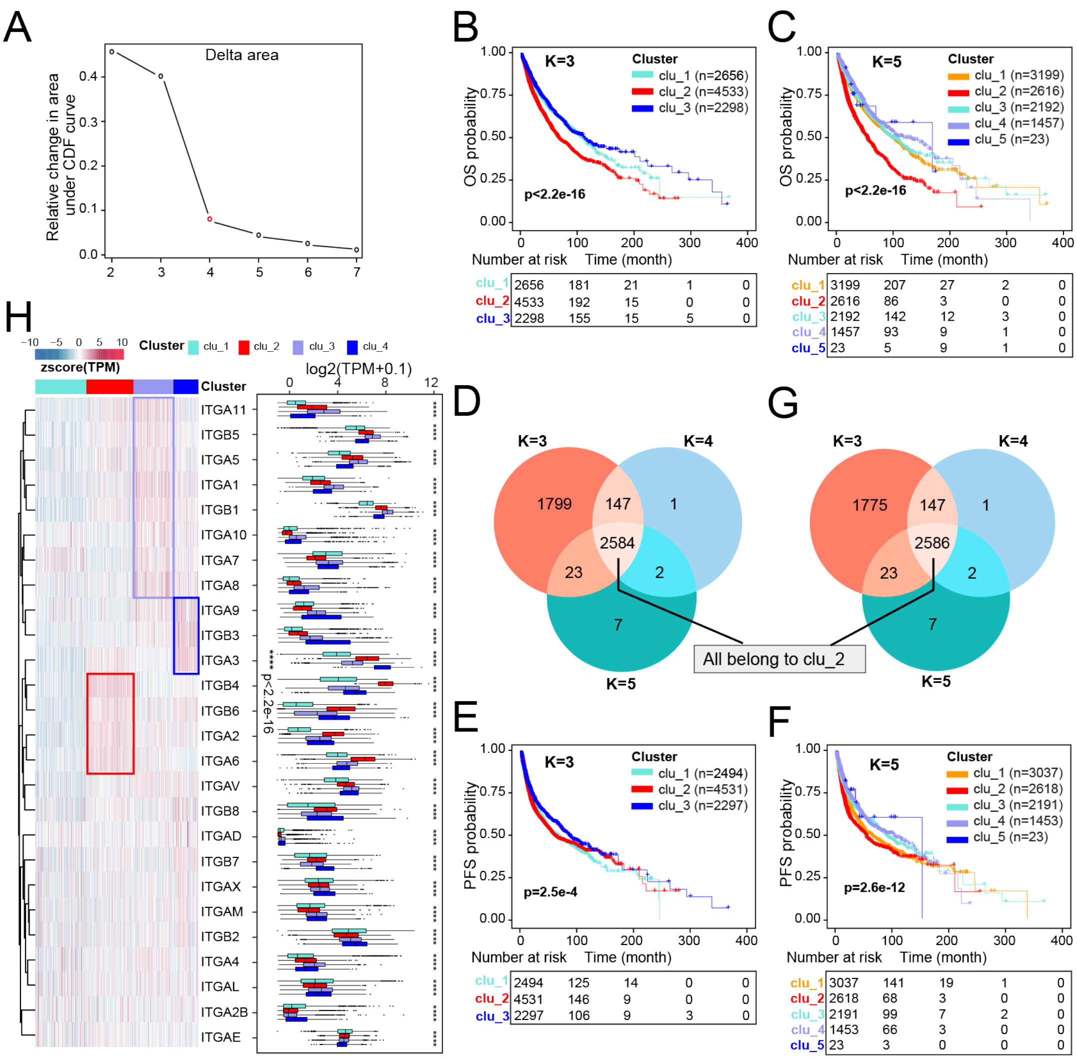


**Fig. S2. Integrin general expression patterns correlate with patient's outcome**

**A.** Unsupervised clustering based on the expression of 26 integrin genes showing k=4 (red point) as the optimal consensus clusters.

**B-C and E-F.** Integrin general expression patterns consistently correlates with OS (**B** and **C**) and PFS (**E** and **F** ) when K=3 (**B** and **E**) and K=5 (**C** and **F**), respectively.

**D and G.** Venn plot showing integrin general expression can convergently identify a subgroup of tumors with worse outcomes (OS and PFS for **D** and **G**, respectively)

**H.** The expression repertoire of 26 integrin genes classifies TCGA tumors into four clusters (termed clu_1/2/3/4; left). Boxplot (right) showing difference in expression of 26 integrin genes among four clusters. Significance is calculated by the Kruskal–Wallis test. Within the boxplots, the center lines represent median values, box edges are 75th and 25th percentiles, and dots denote the outliers.

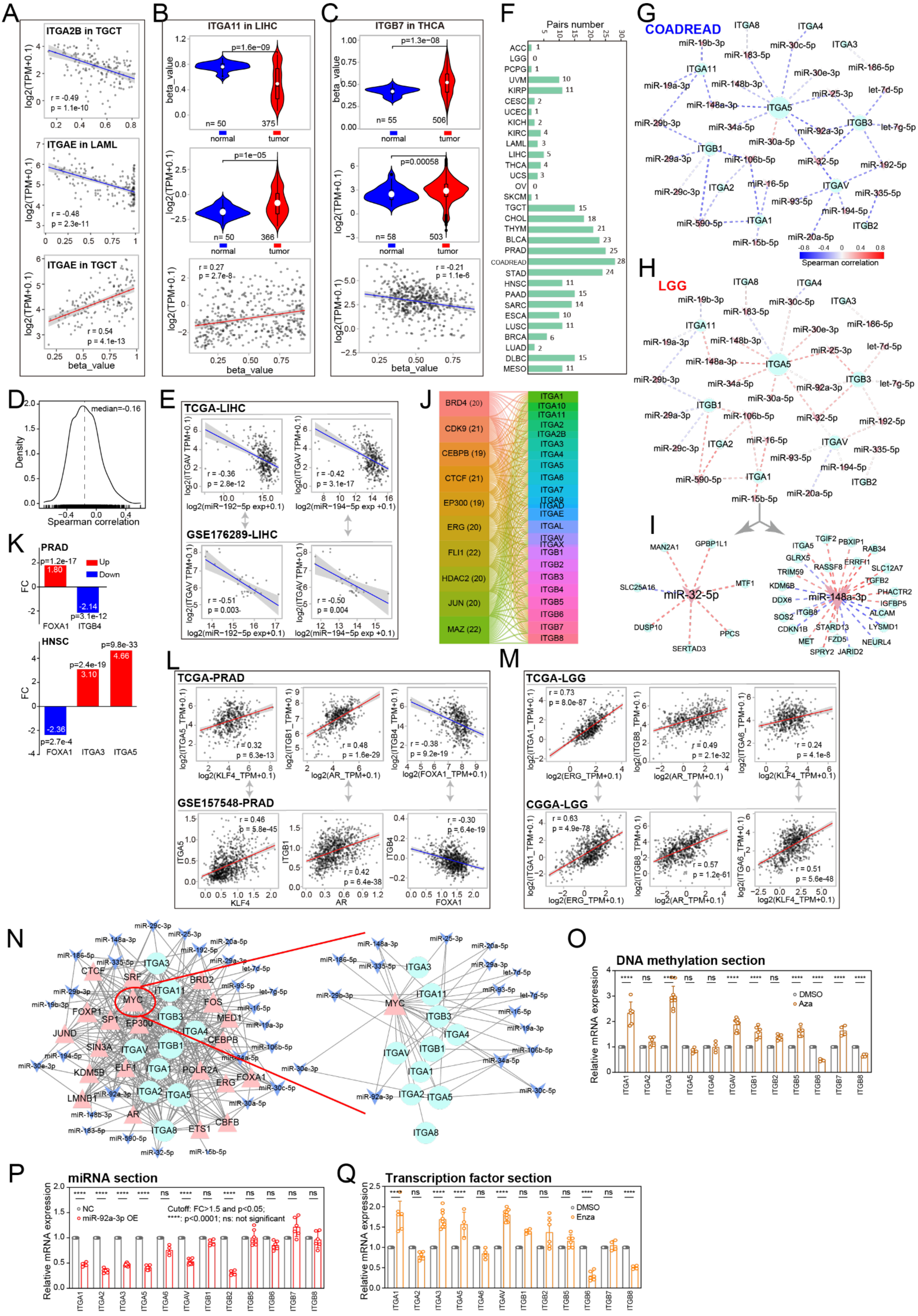
A
ITGA2B in TGCT
r = -0.49
p = 1.1e-10
ITGAE in LAML
r = -0.48
p = 2.3e-11
ITGAE in TGCT
r = 0.54
p = 4.1e-13
log2(TPM+0.1)
beta_value
B
ITGA11 in LIHC
p=1.6e-09
p=1e-05
n= 50
375
366
normal
tumor
r = 0.27
p = 2.7e-8
C
ITGB7 in THCA
p=1.3e-08
p=0.00058
n= 55
506
n= 58
503
r = -0.21
p = 1.1e-6
beta_value
D
median=-0.16
Density
Spearman correlation
E
TCGA-LIHC
r = -0.36
p = 2.8e-12
r = -0.42
p = 3.1e-17
log2(ITGAV TPM+0.1)
log2(miR−192−5p exp+0.1)
log2(miR−194−5p exp +0.1)
GSE176289-LIHC
r = -0.51
p = 0.003
r = -0.50
p = 0.004
F
Pairs number
ACC
LGG
PCPG
UVM
KIRP
CESC
UCEC
KICH
KIRC
LAML
LIHC
THCA
UCS
OV
SKCM
TGCT
CHOL
THYM
BLCA
PRAD
COADREAD
STAD
HNSC
PAAD
SARC
ESCA
LUSC
BRCA
LUAD
DLBC
MESO
G
COADREAD
H
LGG
Spearman correlation
I
miR-32-5p
miR-148a-3p
J
BRD4 (20)
CDK9 (21)
CEBPB (19)
CTCF (21)
EP300 (19)
ERG (20)
FLI1 (22)
HDAC2 (20)
JUN (20)
MAZ (22)
K
PRAD
p=1.2e-17
Up
Down
p=3.1e-12
FOXA1
ITGB4
HNSC
p=9.8e-33
p=2.4e-19
p=2.7e-4
FOXA1
ITGA3
ITGA5
FC
L
TCGA-PRAD
r = 0.32
p = 6.3e-13
r = 0.48
p = 1.6e-29
r = -0.38
p = 9.2e-19
log2(KLF4_TPM+0.1)
log2(AR_TPM+0.1)
log2(FOXA1_TPM+0.1)
GSE157548-PRAD
r = 0.46
p = 5.8e-45
r = 0.42
p = 6.4e-38
r = -0.30
p = 6.4e-19
KLF4
AR
FOXA1
M
TCGA-LGG
r = 0.73
p = 8.0e-87
r = 0.49
p = 2.1e-32
r = 0.24
p = 4.1e-8
log2(ERG_TPM+0.1)
log2(AR_TPM+0.1)
log2(KLF4_TPM+0.1)
CGGA-LGG
r = 0.63
p = 4.9e-78
r = 0.57
p = 1.2e-61
r = 0.51
p = 5.6e-48
N
O
DNA methylation section
Relative mRNA expression
DMSO
Aza
P
miRNA section
NC
miR-92a-3p OE
Cutoff: FC>1.5 and p<0.05;
****: p<0.0001; ns: not significant
Q
Transcription factor section
DMSO
Enza

**Fig. S3. Multiple mechanisms combinedly regulate integrin expression**

**A.** Dot plots showing Spearman correlation between DNA methylation in promoter and expression of integrin genes. Two integrin genes in two cancers are shown.

**B and C.** Inconsistency between expression and methylation level for ITGA11 in LIHC (**B**) and ITGB7 in THCA (**C**). Top and middle: violin plots showing DNA methylation in promoter and expression of integrin gene in tumors and normal tissues, respectively; bottom: dot plots showing Spearman correlation between DNA methylation in promoter and expression of integrin genes. Within the violin plots, the center white dots represent mean values, box edges are 75th and 25th percentiles, and dots denote the outliers.

**D.** Density plots showing correlation coefficient between 40 pairs of miRNAs and integrin genes in TCGA tumors. The median is labeled.

**E**. Validation of the correlation between miRNA and integrin gene in TCGA (top panel) and an independent liver cancer cohort (bottom panel).

**F.** Bar plots showing the number of significant correlated miRNA-integrin pairs across cancers. The cutoff of Spearman $\rho \geq 0.3$ and $p < 0.05$ is considered significant.

**G-I.** Correlation networks of miRNA-integrin pairs in COADREAD (**G**) and LGG (**H**), and miRNA-targets pairs (**I**). The same miRNA-integrin pairs have predominantly negative correlations in COADREAD but positive correlations in LGG. Two miRNAs with positive correlation with part of their targets in LGG are shown as examples. The size of circle or V-shape plots denotes the number of pairs directly associate with this gene.

**J.** Alluvial plot showing the top ten TFs with most integrin targets. Numbers in the parentheses are the number of integrin targets.

**K.** Bar plots showing fold change (FC) of FOXA1 and its integrin targets in PRAD and HNSC tumors compared with normal tissues. P values are labeled.

**L-M.** The correlation between TFs and integrin genes are validated in different cohorts (**K** for PRAD; **L** for LGG). Dot plots show the relationship between individual TF and integrin gene labeled by correlation coefficients and p values.

**N**. A three-way interacting network between miRNA, TFs, and integrin genes (left). A MYC-centered network is shown as an example (right). Lines indicate regulation relationship. The size of circle, triangle, and V-shape plot denotes the number of pairs directly associate with this gene. Circles, integrin genes; triangle, TFs; V-shape plots: miRNAs.

**O-Q**. qPCR analysis of 12 integrin gene expression in prostate cancer LNCaP cells treated with5'-Aza (**O**), miR-92a-3p (**P**), and Enza (**Q**). The cutoff for significance is fold change > 1.5 and p-value < 0.05.

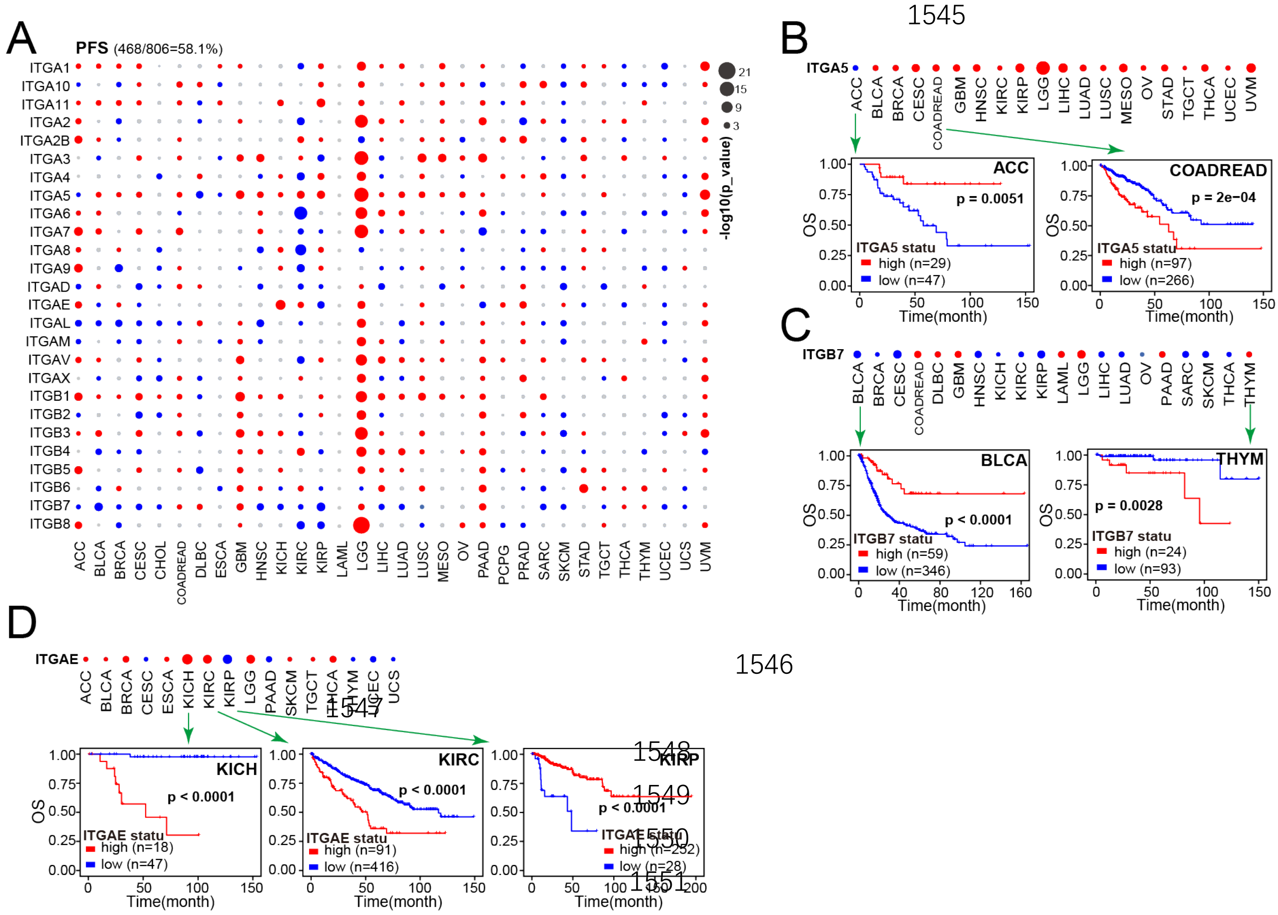


**Fig. S4. Heterogeneity of clinical relevance of integrin genes in a cancer-depend manner.**

**A.** Summary of progression free survival (PFS) time analysis results of integrin genes across cancers. Red, blue and grey dots indicate worse, better and unsignificant results, respectively. The size of dots indicates -log2(p_values). Data in the parentheses (top left) denotes the rate of dysregulation associated with PFS.

**B-D**. Three examples (**B** for ITGA5, **C** for ITGB7, and **D** for ITGAE) of cancer-specific associations between indicated integrin gene expression and overall survival (OS).

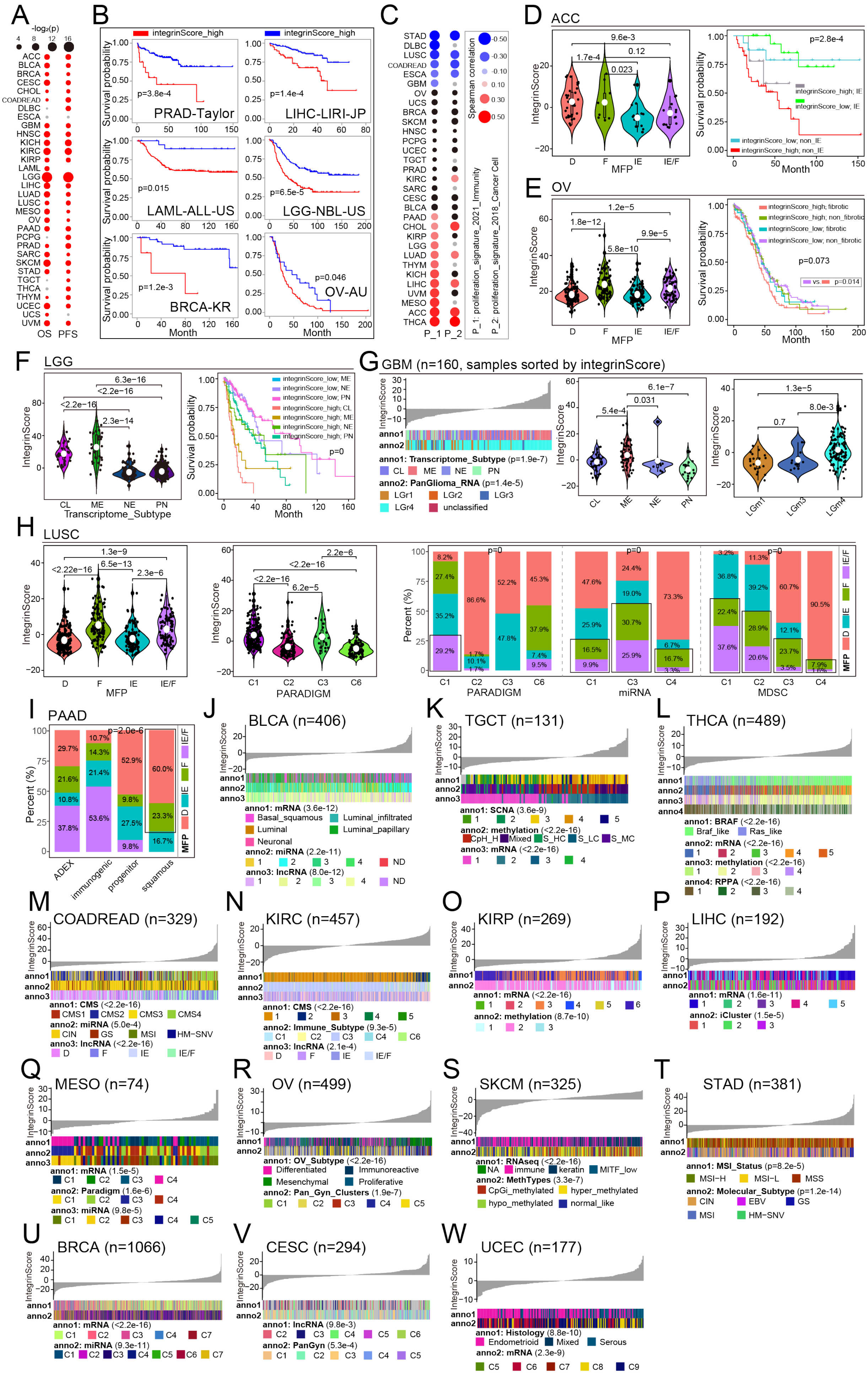
A
-log₂(p)
4 8 12 16
ACC BLCA BRCA CESC CHOL COADREAD DLBC ESCA GBM HNSC KICH KIRC KIRP LAML LGG LIHC LUAD LUSC MESO OV PAAD PCPG PRAD SARC SKCM STAD TGCT THCA THYM UCEC UCS UVM
OS PFS
B
integrinScore_high
integrinScore_high
Survival probability
p=3.8e-4
PRAD-Taylor
p=1.4e-4
LIHC-LIRI-JP
p=0.015
LAML-ALL-US
p=6.5e-5
LGG-NBL-US
p=1.2e-3
BRCA-KR
p=0.046
OV-AU
Month
C
STAD DLBC LUSC COADREAD ESCA GBM OV UCS BRCA SKCM HNSC PCPG UCEC TGCT PRAD KIRC SARC CESC BLCA PAAD CHOL KIRP LGG LUAD THYM KICH LIHC UVM MESO ACC THCA
P_1 P_2
Spearman correlation
-0.50 -0.30 -0.10 0.10 0.30 0.50
P_1: proliferation_signature_2021_Immunity
P_2: proliferation_signature_2018_Cancer Cell
D ACC
9.6e-3
1.7e-4
0.12
0.023
IntegrinScore
MFP
D F IE IE/F
p=2.8e-4
integrinScore_high; IE
integrinScore_low; IE
integrinScore_low; non_IE
integrinScore_high; non_IE
Survival probability
Month
E OV
1.2e-5
1.8e-12
9.9e-5
5.8e-10
IntegrinScore
MFP
integrinScore_high; fibrotic
integrinScore_high; non_fibrotic
integrinScore_low; fibrotic
integrinScore_low; non_fibrotic
p=0.073
vs p=0.014
Survival probability
Month
F LGG
6.3e-16
<2.2e-16
<2.2e-16
2.3e-14
IntegrinScore
CL ME NE PN
Transcriptome_Subtype
integrinScore_low; ME
integrinScore_low; NE
integrinScore_low; PN
integrinScore_high; CL
integrinScore_high; ME
integrinScore_high; NE
integrinScore_high; PN
p=0
Survival probability
Month
G GBM (n=160, samples sorted by integrinScore)
IntegrinScore
anno1
anno2
anno1: Transcriptome_Subtype (p=1.9e-7)
CL ME NE PN
anno2: PanGlioma_RNA (p=1.4e-5)
LGr1 LGr2 LGr3 LGr4 unclassified
6.1e-7
5.4e-4
0.031
1.3e-5
8.0e-3
0.7
LGm1 LGm3 LGm4
H LUSC
1.3e-9
<2.22e-16
6.5e-13
2.3e-6
IntegrinScore
MFP
2.2e-6
<2.2e-16
<2.2e-16
6.2e-5
C1 C2 C3 C6
PARADIGM
Percent (%)
p=0
miRNA
MDSC
IE/F F IE D MFP
I PAAD
p=2.0e-6
Percent (%)
ADEX immunogenic progenitor squamous
J BLCA (n=406)
anno1: mRNA (3.6e-12)
Basal_squamous Luminal_infiltrated Luminal Luminal_papillary Neuronal
anno2: miRNA (2.2e-11)
anno3: lncRNA (8.0e-12)
K TGCT (n=131)
anno1: SCNA (3.6e-9)
anno2: methylation (<2.2e-16)
CpH_H Mixed S_HC S_LC S_MC
anno3: mRNA (<2.2e-16)
L THCA (n=489)
anno1: BRAF (<2.2e-16)
Braf_like Ras_like
anno2: mRNA (<2.2e-16)
anno3: methylation (<2.2e-16)
anno4: RPPA (<2.2e-16)
M COADREAD (n=329)
anno1: CMS (<2.2e-16)
CMS1 CMS2 CMS3 CMS4
anno2: miRNA (5.0e-4)
CIN GS MSI HM-SNV
anno3: lncRNA (<2.2e-16)
D F IE IE/F
N KIRC (n=457)
anno1: CMS (<2.2e-16)
anno2: Immune_Subtype (9.3e-5)
anno3: lncRNA (2.1e-4)
O KIRP (n=269)
anno1: mRNA (<2.2e-16)
anno2: methylation (8.7e-10)
P LIHC (n=192)
anno1: mRNA (1.6e-11)
anno2: iCluster (1.5e-5)
Q MESO (n=74)
anno1: mRNA (1.5e-5)
anno2: Paradigm (1.6e-6)
anno3: miRNA (9.8e-5)
R OV (n=499)
anno1: OV_Subtype (<2.2e-16)
Differentiated Immunoreactive Mesenchymal Proliferative
anno2: Pan_Gyn_Clusters (1.9e-7)
S SKCM (n=325)
anno1: RNAseq (<2.2e-16)
NA immune keratin MITF_low
anno2: MethTypes (3.3e-7)
CpGi_methylated hyper_methylated hypo_methylated normal_like
T STAD (n=381)
anno1: MSI_Status (p=8.2e-5)
MSI-H MSI-L MSS
anno2: Molecular_Subtype (p=1.2e-14)
CIN EBV GS MSI HM-SNV
U BRCA (n=1066)
anno1: mRNA (<2.2e-16)
anno2: miRNA (9.3e-11)
V CESC (n=294)
anno1: lncRNA (9.8e-3)
anno2: PanGyn (5.3e-4)
W UCEC (n=177)
anno1: Histology (8.8e-10)
Endometrioid Mixed Serous
anno2: mRNA (2.3e-9)

**Fig. S5. IntegrinScore correlates with outcomes and known molecular subtypes in distinct cancers**

**A.** Summary of overall survival (OS) and progression free survival (PFS) time analysis results of integrinScore across cancers. Red and grey dots indicate worse and unsignificant results, respectively. The size of dots indicates $-\log_2$(p_values).

**B.** Kaplan-Meier plots showing integrinScore consistently correlates with worse outcome in distinct cohorts.

**C.** Spearman correlation of integrinScore with two proliferation related signatures. Blue, red, black and grey dots denote positive, negative and unsignificant correlations, and absent values, respectively. The size of dots indicates the absolute correlation coefficient.

**D-F.** Left: violin plots showing integrinScore among four TME subtypes in ACC (**D**), OV (**E**) and LGG (**F**); right: Kaplan-Meier plots showing integrinScore combined with known subtypes can further identify more aggressive tumors.

**G.** Left: an overview of the association between known molecular subtypes and integrinScore in GBM. Middle and right: violin plots showing integrinScore among distinct molecular subtypes.

**H.** Left and middle: violin plots showing integrinScore among distinct molecular subtypes in LUSC. Right: Comparison of the percentage of four TME subtypes among distinct subgroups of PARADIGM, miRNA and MDSC. Chi-square test is used, and so as to **I**.

**I.** Comparison of the percentage of four TME subtypes among subgroups in PAAD.

**J-W**. An overview of the association between known molecular subtypes and integrinScore across in BLCA (**J**), TGCT (**K**), THCA (**L**), COADREAD (**M**), KIRC (**N**), KIRP (**O**), LIHC (**P**), MESO (**Q**), OV (**R**), SKCM (**S**), STAD (**T**), BRCA (**U**), CESC (**V**), and UCEC (**W**). 挑 2 个加一点细节，正文里面没有说，legend 要描述一下。

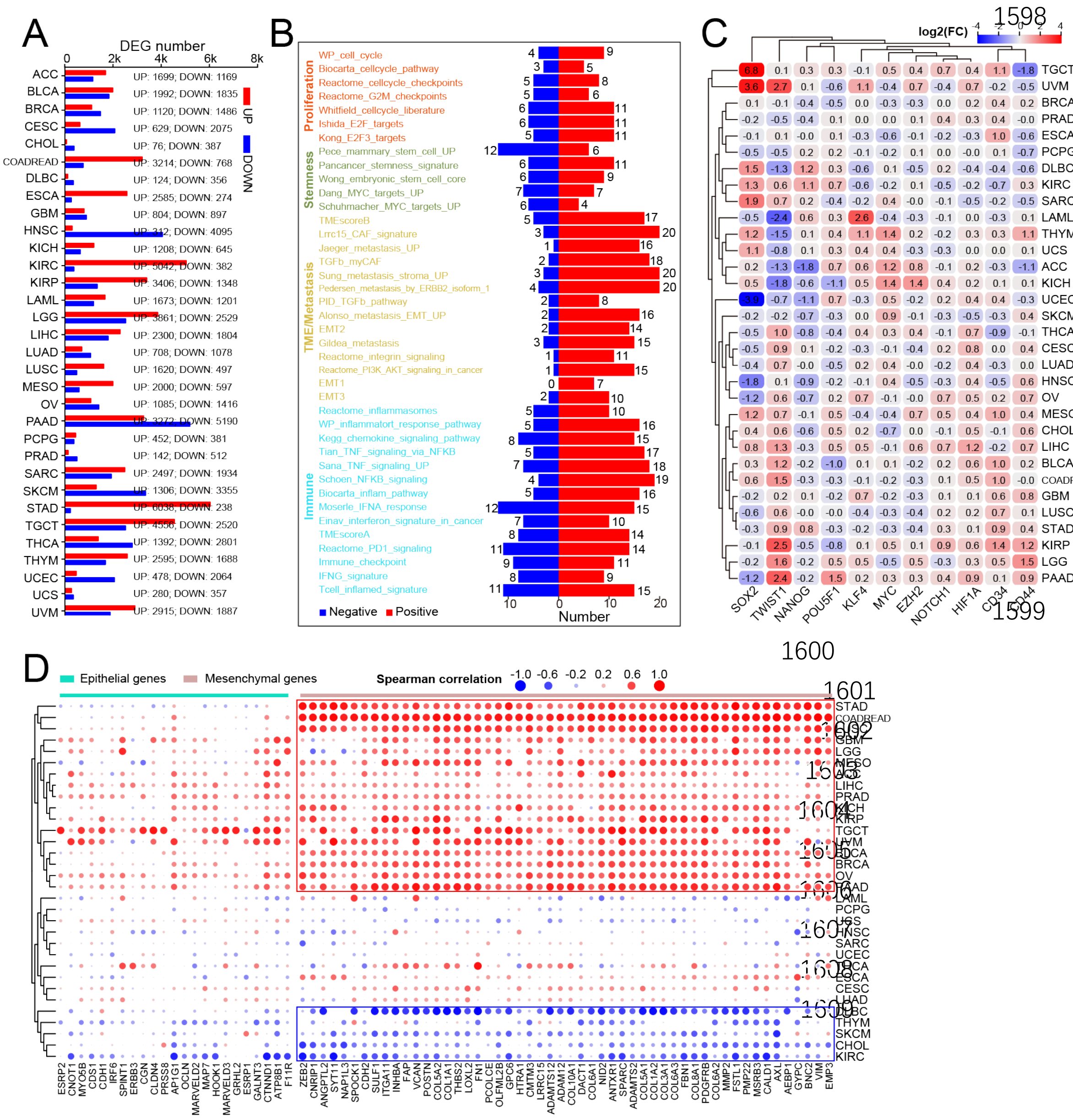


**Fig. S6. IntegrinScore correlates with stemness and immune markers in cancer**

**A.** Bar plots showing the number of differentially expressed genes (DEGs) between integrinScore high and low groups across cancers.

**B.** Bar plots showing the number of cancer types in which each signature significantly enriches. Four types of signatures are labeled with different colors.

**C.** Heatmap showing foldchange (FC) of stemness markers between integrinScore-high and low groups across cancers. Red and bule colors denote up- and down-regulation of genes. FCs are labeled in boxes.

**D.** Spearman correlation of integrinScore with expression of epithelial and mesenchymal markers. Blue and red dots denote positive and negative correlations, respectively. The size of dots indicates the absolute correlation coefficient.

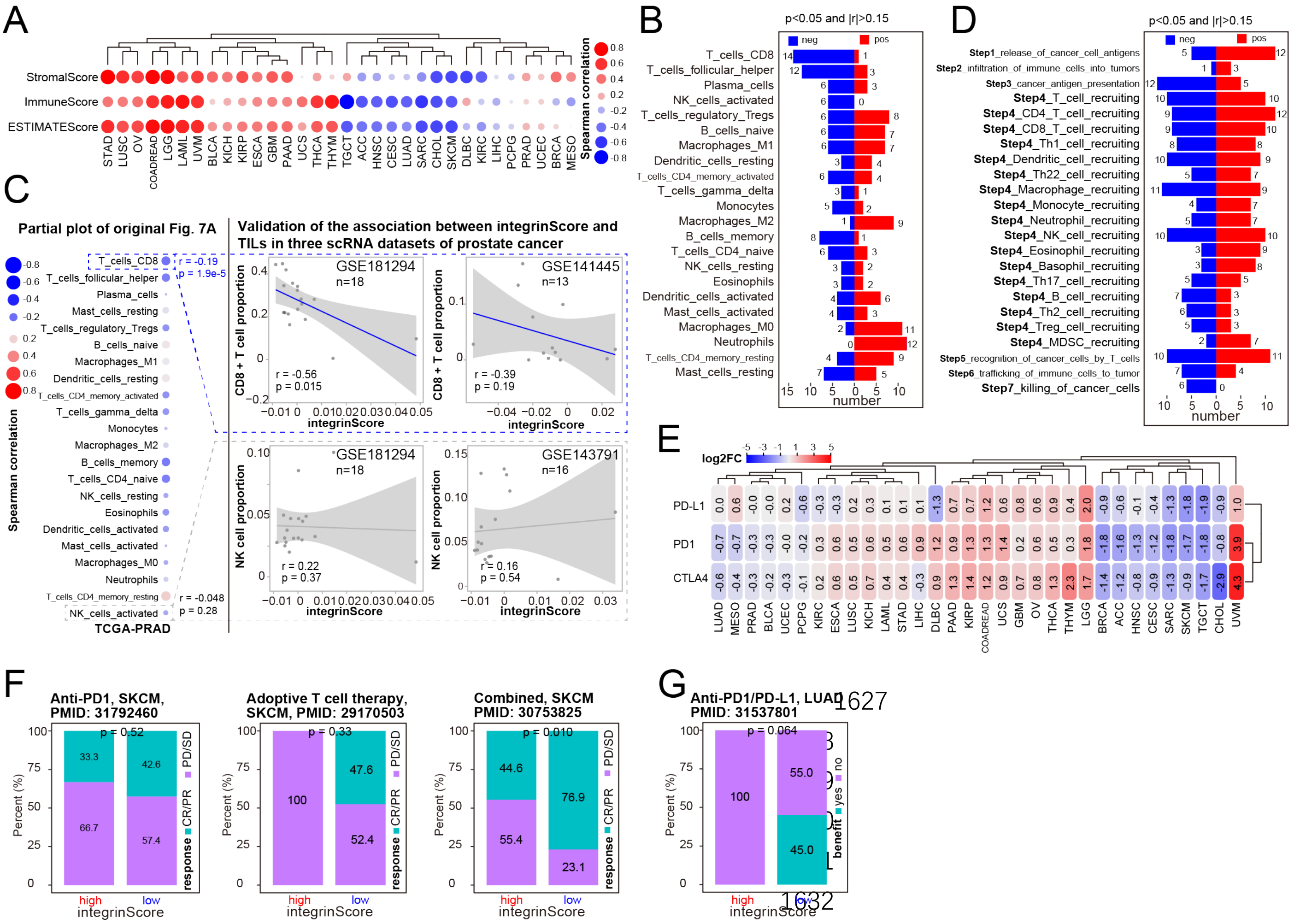


**Fig. S7. IntegrinScore predicts patient responses to immune therapy**

**A.** Spearman correlation of integrinScore with immune landscape score across cancers. Blue and red dots denote positive and negative correlations, respectively. The size of dots indicates the absolute correlation coefficient.

**B**. Bar plots showing the number of cancer types in which integrinScore significantly correlates with immune cell abundance. The cutoff is labeled on the top.

**C**. The correlation between integrinScore and the proportion of CD8+ T cell and NK cell in three single-cell prostate cancer (PCa) datasets. Three PRAD scRNA-seq datasets (GSE181294, GSE141445 and GSE143791) were downloaded from GEO database. After standard quality control, we annotated the cell identity for each subpopulation by commonly-used lineage markers. The integrinScore was deduced based on average transcriptome by treating all single-cells as a whole for a sample. Finally, we calculated the correlation between integrinScore and TIL proportions by Spearman correlation analysis.

**D.** Bar plots showing the number of cancer types in which integrinScore significantly correlates with immune steps. The cutoff is labeled on the top.

**E.** Heatmap showing fold change (FC) of three ICB markers between integrinScore-high and low groups across cancers. Red and bule colors denote up- and down-regulation of genes. FCs are labeled in boxes.

**F and G.** Comparison of the percentage of patients response to immune therapies between integrinScore high and low group in SKCM (**F**) and LUAD (**G**). Chi-square test is used.

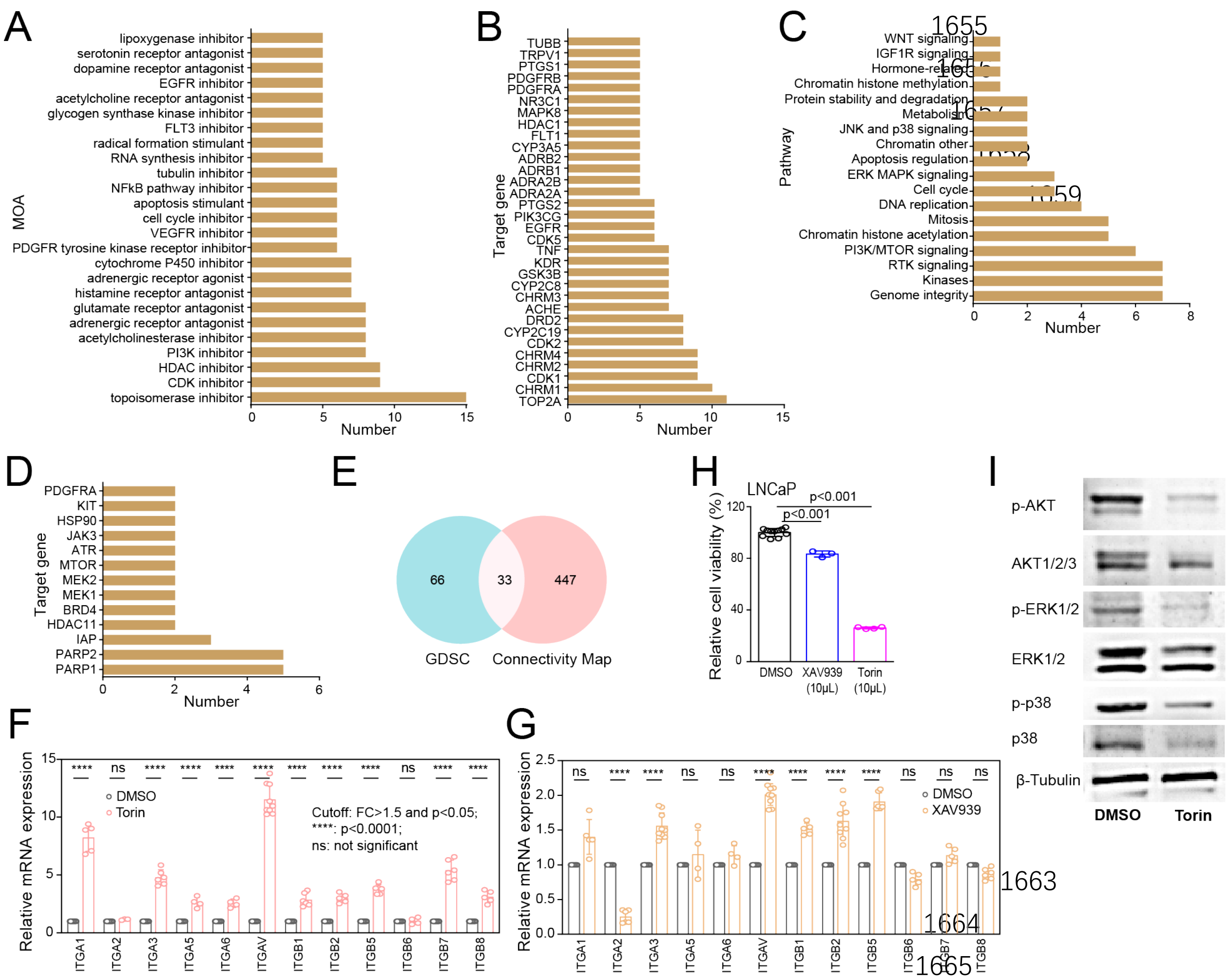


**Fig. S8. Pathways and target genes of drugs correlated with integrinScore**

**A and B.** Bar plots showing top mechanisms of action (MOA; **A**) and target genes (**B**) of compounds correlated with integrinScore by the CMap analysis.

**C and D.** Bar plots showing top pathways (**C**) and target genes (**D**) of compounds correlated with integrinScore in the GDSC database.

**E.** Venn plot showing the overlapping of target genes based on the CMap and GDSC database.

**F** and **G**. qPCR analysis of 12 integrin gene expression in LNCaP cells treated with Torin2 (**F**) and XAV939 (**G**).

**H**. Cell viability (MTT assay) in LNCaP cells treated with Torin and XAV939 for 3~4 days.

**I**. Western blot analysis of indicated mitotic pathways in LNCaP cells treated with Torin (50 nM for 3 days).
